# The DNA binding protein BCL6 regulates NFκB-controlled endothelial inflammatory gene expression

**DOI:** 10.1101/2022.11.03.514942

**Authors:** Adriana Franco Acevedo, Julia J. Mack, Nicole M Valenzuela

## Abstract

**Background:** NFκB drives acute vascular inflammation by activating gene expression programs in endothelial cells to promote leukocyte recruitment. Numerous negative feedback regulators of NFκB activation have been defined that promote resolution of inflammation. However, the identities of endogenous suppressors of NFκB transcription are less clear. In macrophages, the transcriptional repressor BCL6 was shown to substantially overlap with NFκB-driven genes and influence the response to LPS. We identified that the DNA binding protein BCL6 was expressed in endothelial cells. Although the role of BCL6 in adaptive immune cells has been characterized, how BCL6 modifies transcription in endothelial cells has not been studied.

**Objective:** Based on prior knowledge that BCL6 represses part of the LPS-induced transcriptome in macrophages, we asked whether BCL6 regulated endothelial pro-inflammatory state by direct interaction with NFκB.

**Methods:** We analyzed public datasets of RNA and ChIP-Seq, probed BCL6 expression in human tissue, and tested BCL6 knockdown, overexpression and pharmacological manipulation on TNFα induced gene expression *in vitro* using human primary endothelium isolated from the heart.

**Results:** We demonstrate that the DNA binding protein BCL6 is basally expressed in the endothelium, with chromatin marks reflective of a superenhancer, and is particularly enriched in aortic endothelial cells (ECs) compared with ECs from other organs. Although basal expression was relatively low, BCL6 was rapidly upregulated in cardiac endothelium stimulated with TNFα, through direct action of NFκB. The BCL6 consensus DNA binding motif overlaps with that of NFκB. BCL6 target genes included endothelial pro-inflammatory chemokines and adhesion molecules, as well as NFκB-related genes themselves. BCL6 knockdown and the degrading BCL6 inhibitor BI-3802 augmented the endothelial cell response to TNFα. Surprisingly, antagonism of the BTB domain of BCL6 with small molecules 79-6, FX1 or BI-3812, blocked leukocyte adherence and accordingly suppressed both NFκB transcriptional activity as well as the expression of many genes in response to TNFα. Lastly, we show that HDAC activity is increased by TNFα, and can be reduced in the presence of BTB domain inhibitors.

**Conclusions:** Our results demonstrate that BCL6 is a repressor of NFκB-driven gene expression and inflammation in cardiac endothelial cells. These findings indicate that targeting of BCL6 may enhance vascular inflammation resolution.

## Introduction

The resolution of inflammation limits the duration and magnitude of an inflammatory response and actively accelerates the return to tissue homeostasis[1]. The vascular endothelial cell (EC) lining the lumen of blood vessels is capable of initiating local inflammatory responses through leukocyte recruitment, and modifying the adaptive immune system by costimulatory and antigen presentation functions [1, 2]. Yet, the mechanisms essential for endothelial resilience or termination of peripheral inflammation are incompletely defined.

Endothelial cells are central to the initiation and resolution of vascular inflammation in autoimmune vasculitis and transplant rejection, but are rarely a direct target of therapeutic intervention. It is increasingly recognized that endothelial cells exhibit remarkable tissue-specific diversity in transcriptome, phenotype and function [3]. However, the divergent inflammatory responses of endothelial cells from tissue-specific vascular beds remain incompletely characterized, and the underlying mechanisms are poorly understood. In a recent study we investigated whether tissue origin of endothelial cells (EC) influenced the magnitude and kinetics of adhesion molecule expression, chemokine patterns and resultant leukocyte adherence [4]. In particular, we found that cardiac endothelium exhibits a greater response to inflammatory stimuli compared with endothelial cells from liver and kidney. Our results showed twenty-seven immune-associated genes that were differentially expressed in untreated EC. Among them, *BCL6* (ZBTB27) [4] was particularly enriched in cardiac endothelium and correlated with more intense endothelial inflammatory responses.

BCL6 is a DNA binding protein originally identified as an oncogene in lymphoid malignancy [5, 6] and that subsequently was recognized as a lineage-defining factor indispensable for T follicular helper cell development [7]. Specifically, BCL6 functions as a transcriptional repressor important in germinal center responses and Tfh cell differentiation [8, 9]. BCL6 is composed of a sequence-specific DNA binding zinc finger domain; a linker region; and a BTB domain through which it complexes with corepressors such as BCOR, and SMRT:NCOR [10, 11]. These corepressor associations enable epigenetic regulation of target genes through recruitment of class I histone deacetylases (HDAC), including HDAC3 [12]. BCL6 also recruits class II HDACs such as HDAC7 [13]. In murine macrophages, BCL6 inhibition or deficiency significantly alter chemokine gene expression and perturb inflammatory gene modules in response to LPS [11, 14]. Here, BCL6 overlapped with the NFκB cistrome, and regulated approximately one fourth of NFκB-driven genes.

Outside the immune system, BCL6 exhibits variable tissue distribution, with high transcript in the heart and lung [15]. Systemic knockout of *Bcl6* in mice caused not only germinal center and Th17 defects but also massive myocarditis and pulmonary inflammation, which developed between 4-6 weeks of age [16, 17]. Importantly, myocarditis was attributable to cardiac-intrinsic *Bcl6* rather than to augmented responses in the hematopoietic compartment [18].

Several lines of evidence imply that BCL6 is important in vascular health and disease. First, BCL6 is significantly associated with cardiovascular disease in GWAS studies [19]. BCL6 expression is downregulated in other forms of human cardiovascular disease including cardiomyopathies [20, 21]; obliterative pulmonary hypertension [22, 23]; and heart failure [24]. Its suppression in the mouse aorta was associated with elevated inflammatory genes in atherosclerosis [25]. Along with endothelial-specific ETS2, BCL6 interacted with the safeguard transcription factor TWIST1 [26], the latter of which is implicated as a causal gene in human vascular disease [27] particularly functionally linked with endothelial-initiated atherosclerosis [28].

BCL6 was recently identified as a key transcription factor governing blood over lymphatic endothelial cell identity [29]. However, only a few studies have investigated the function of BCL6 in the blood endothelium. Both BCL6 and its corepressor BCOR were induced in EC by tumor-conditioned medium, VEGF and angiopoietin-2, and loss of BCL6 reduced endothelial proliferation while augmenting angiogenic sprouting through DLL4 [30]. In another study, PPARδ agonists suppressed VCAM-1 expression by increasing BCL6 binding to the *VCAM1* promoter [31]. However, BCL6 does not otherwise have a characterized role in the regulation of endothelial cell inflammatory response.

A major mechanism of transcriptional repression mediated by BCL6 is to associate with corepressors that recruit histone deacetylases. BCL6 known interactors include corepressor complex BCOR, NCOR2/SMRT, HDAC; Jun family, JNK interacting proteins and Sp1 transcription factors; IRF4; p300; and NFκB repressing factor; and the potential regulatory network through secondary interactions is very wide [32]. In B cells, BCL6 preferentially localized to less accessible chromatin regions and negatively correlated with transcriptional activity of nearby genes [33]. BCL6 cooperates with NCOR/SMRT in immune cells to recruit HDAC3, decrease H3K27Ac and repress transcription [10, 12, 34–36]. In macrophages, the BCL6 cistrome partially overlaps with that of NFκB, where its deletion or inhibition augments a program of chemokines and other pro-inflammatory mediators in the response to LPS and in atherogenesis [11, 14].

Because BCL6 is an oncogenic target in lymphoma, several small molecule inhibitors have been developed to either degrade the protein or antagonize its function. BCL6 degrading molecules, such as BI-3802 [37] [38] or proteolysis-targeting chimeras (PROTACs) [39], cause rapid degradation of BCL6 protein.

Non-degrading inhibitors, such as 79-6, and its high potency derivative FX1, block the BTB domain and prevent association of BCL6 with corepressors NCOR/SMRT [40, 41]. This reactivated a number of BCL6-repressed target genes in lymphoma cells [42]. Another molecule, BI-3812, bound to the BTB domain, reduced complex formation with NCOR1, and substantially altered gene expression in BCL6-dependent B cell lymphoma lines [37]. Similarly, the peptidomimetics RI-BPI [43, 44] and GSK137 [45] bind the lateral groove of BCL6 and dissociate corepressors. Not only did FX1 prevent germinal center formation *in vivo* [46] but it also showed efficacy in reducing experimental end-stage organ damage in autoimmune nephritis [47] and sepsis [48].

In this study, we posited that BCL6 acts as a mediator of inflammation resolution, by cooperatively modulating the initiation, magnitude and termination of endothelial cell immune responses via NFκB. First, we show that endothelial cells broadly express BCL6, although at a lower basal level than immune cells. Then, we demonstrate that BCL6 expression among endothelial cell types is particularly enriched in the heart and lung compared with other organs, in both mouse and human. Moreover, in endothelium, the *BCL6* gene is proximally downstream of a superenhancer with high chromatin accessibility. In addition to basal expression, we provide evidence that BCL6 is inducible in endothelium in response to many stimuli, including the pro-inflammatory cytokine TNFα, and that RELA/NFκB regulates its expression. Lastly, we show that knockdown, overexpression or treatment with pharmacological agents that target BCL6 significantly alter the endothelial response to TNFα, predominantly by NFκB-driven targets, both *in vitro* and *in vivo*. Surprisingly, BTB domain inhibitors and overexpression suppress, while BCL6 depletion potentiates, inflammatory activation of the endothelium by TNFα, including chemokine production, adhesion molecule expression and binding of leukocytes. Our findings describe a novel pathway of BCL6-regulated endothelial cell activation, representing an untapped therapeutic target for resolving vascular inflammation.

## Materials and Methods

### Ethics Statement

Use of human endothelial and peripheral blood cells was approved by the UCLA Institutional Review Board (IRB17-00477). IRB was waived for the use of deidentified human tissue from the UCLA Translational Pathology Core Laboratory.

### Human Tissue

Archived FFPE human normal spleen, heart, lung, kidney and liver tissue were procured from the UCLA Pathology Biorepository. Samples were fully anonymized; race, ethnicity sex and age are not available. Tissues were stained for CD31 (vendor), ERG (vendor), BCL6 (vendor), and DAPI immunofluorescence by the UCLA Translational Pathology Core Laboratory. Slides were digitally scanned by XX and analyzed in XX.

### Cells

Primary human endothelium from the aorta (HAEC), coronary artery (HCAEC), cardiac microvasculature (HCMVEC), pulmonary artery (HPAEC), lung microvasculature (HLMVEC) or liver sinusoids (HLSEC) were purchased from ATCC, PromoCell and Lonza. NFKB-TIME, an immortalized human microvascular endothelial cell line stably transfected with NFκB-responsive pNLuc, was obtained from ATCC and cultured in complete ECM (ATCC). Cells were cultured in complete endothelial cell growth medium (PromoCell, ATCC) and stimulated in M199 + 10% heat-inactivated fetal bovine serum (FBS). After receipt and initial subculturing, primary and immortalized endothelial cells were authenticated by staining for CD31, and upregulation of VCAM-1 expression after 4hr of TNFα stimulation, before use in assays. Peripheral blood mononuclear cells (PBMC) were prepared from the whole blood of healthy volunteers by Ficoll-Hypaque density centrifugation, or purchased from ATCC.

### Reagents

The following inhibitors were obtained from commercial sources: BI-3812, 79-6 and BI-3802 were obtained from Selleck Chemical. The NFκB/NLRP3 inhibitor Bay11-7028 was obtained from Invivogen. Human TNFα and FX1 were purchased from Sigma. The broad HDAC inhibitor Trichostatin A (TSA), class I HDAC inhibitor CI994 and histone acetyltransferase inhibitor II (HATi) were purchased from Selleck Chemical.

### siRNA Knockdown

Endothelial cells were seeded at 50-70% confluence. The next day, medium was replaced with 1μM GAPDH or BCL6 siRNA in Accell Delivery Medium (Dharmacon/Horizon). Cells were supplemented with fresh normal growth medium 24-48hr after siRNA addition, and cultured for a total of 24-96hr before gene expression measurements.

### Lentiviral Transduction for BCL6 overexpression and knockdown

For overexpression, endothelial cells were seeded at 40-70% confluence, then infected with third generation lentiviral particles encoding the BCL6 open reading frame (pLenti-BCL6 ORF-mGFP) or GFP alone (Origene). For knockdown, endothelium was infected with lentivirus encoding shRNA against BCL6 (pLenti-BCL6 shRNA) or GAPDH (Horizon). Cells were transduced with lentiviral particles in complete medium supplemented with 8μg/mL overnight. Within 24hr, medium was refreshed and cells were incubated for another 24hr to confluence, and reporter GFP expression was verified, before expanding or use in experiments.

### Nanostring

Nanostring assays were performed by the UCLA Center for Systems Biomedicine. For transcript changes, endothelial monolayers were detached with trypsin, pelleted by centrifugation, and resuspended in RLT Buffer at 6,500 cells per μL (Qiagen). mRNA counts were measured by Nanostring (Human Immunology Panel v2.0, Nanostring Technologies) and analyzed in NCounter software. mRNA counts were normalized against internal and housekeeping controls. Normalized counts ≥250 were considered positive, and genes were considered changed if the counts differed by ±50% (i.e. 1.5-fold) or more compared with baseline.

### RNA-Sequencing

RNA Sequencing was performed by the UCLA Technology Center for Genomics and Bioinformatics. For transcript changes, endothelial monolayers from 6 well plates were detached with trypsin, pelleted by centrifugation, and resuspended in 350μL of RLT Buffer (Qiagen). RNA was prepared by RNeasy kit (Qiagen). Libraries for RNA-Seq were prepared with KAPA Stranded mRNA-Seq Kit. RNA-Sequencing workflow consists of mRNA enrichment and fragmentation, first strand cDNA synthesis using random priming followed by second strand synthesis converting cDNA:RNA hybrid to double-stranded cDNA (dscDNA), and incorporates dUTP into the second cDNA strand. cDNA generation is followed by end repair to generate blunt ends, A-tailing, adaptor ligation and PCR amplification. Different adaptors were used for multiplexing samples in one lane. Sequencing was performed on Illumina HiSeq 3000 for SE 1×65 run. Data quality check was done on Illumina SAV. Demultiplexing was performed with Illumina Bcl2fastq v2.19.1.403 software. The Partek flow was used for the data analysis. The alignment was performed using STAR with human reference genome hg38. The hg38 - Ensembl Transcripts release 104 gtf was used for gene feature annotation. For normalization of transcripts counts, 1.0E-4 was added to all values followed normalization by counts per million (CPM) in Partek. Alternatively, 1.0E-4 was added to all values followed normalization by TMM normalization method in Partek.

### NFκB Reporter Assay

NFKB-TIME is an hTERT-immortalized endothelial cell line stably transfected with linearized pNL3.2.NF-κB-RE[NlucP/NF-κB-RE/Hygro] plasmid. NFKB-TIME cells express NanoLuc luciferase with a C-terminal PEST domain, temporally responsive to NFκB transcriptional activity. The luciferase reporter cell line NFKB-TIME was cultured to confluence on 0.2% gelatin in a 96 well plate, then pre-treated with inhibitors where indicated and stimulated with cytokine in M199+10% FBS. At the end of stimulation, conditioned medium was removed and replaced with 50-100μL of fresh, room temperature M199+10% FBS. An equal volume of pre-mixed NanoGlo luciferase substrate in NanoGlo Luciferase Assay (Promega) lysis buffer was added to each well. Relative luminescence units (RLU) were read on a Cytation5 microplate imaging reader (Biotek) after 5min but within 3hr of substrate addition.

### Secreted chemokine/cytokine protein measurements

Conditioned medium from endothelial cell cultures was retained and stored at -20⁰C. Supernatants were assayed for secreted chemokines and cytokines using the Milliplex 38-plex Human Cytokine/Chemokine Assay (Millipore Sigma) according to the manufacturer’s protocol. Beads were acquired on a Luminex 100/200 instrument. Luminex assays were performed by the UCLA Immune Assessment Core.

### Intracellular signaling protein measurements

Endothelial cells were cultured to confluence in a 6 well tissue culture plate. Cells were pre-treated with inhibitors and stimulated with cytokine. After stimulation, medium was collected and cells were washed once with ice cold cell wash buffer, then lysed in 250μL of MILLIPLEX Lysis Buffer. Lysates were assayed for phospho proteins using the Milliplex Cell Signaling kit (NFκB Magnetic Bead Signaling 6-plex, Millipore Sigma) for detection of NFκB pSer536, IκBα pSer32, and IKKα/β pSer177/181) according to the manufacturer’s protocol. Beads were acquired on a Luminex 100/200 instrument. Luminex assays were performed by the UCLA Immune Assessment Core.

### Lumit protein measurements

Total and phosphorylated NFκB were quantitated by Lumit Immunoassay Cellular System (Promega W1220) according to the manufacturer’s protocol. Briefly, endothelial cells were cultured to confluence in a 96 well black plate, then pre-treated with inhibitors where indicated and stimulated with cytokine in M199+10% FBS. At the end of stimulation, cells were lysed with vendor lysis buffer with Halt Protease Inhibitor Cocktail (ThermoFisher #78430) for 20min on a shaker. After lysis, primary antibody pairs against NFκB total (rabbit-anti-NFκB p65 total, Cell Signaling Technology #8242; mouse-anti-NFκB p65 total, Cell Signaling Technology #6956); NFκB phospho (rabbit-anti-NFκB p65 total, Cell Signaling Technology #8242; mouse anti-NFκB p65 p-Ser536, Cell Signaling Technology #13346), mixed with Lumit BiT Set 2 (anti-mouse Ab-SmBiT, anti-rabbit Ab-LgBiT, Promega W1220) and added for 90min. Then, Lumit substrate was added and luminescence was read on a Cytation5 microplate reader.

### Immunofluorescence

Endothelial cells were cultured to confluence in 0.2% gelatin-coated, glass-bottom tissue culture plates, and stimulated. After stimulation, medium was removed, cells were washed once with PBS and fixed for 10min in 4% PFA in neutral buffer. Cells were washed once more with PBS and then stained for ZO-1 (Invitrogen #MA3-39100-A488), NFκB p65 (Cell Signaling #8242), BCL6 (Cell Signaling #14895T), and DAPI (Cell Signaling #4083), then stained with species fluorescent secondary antibodies. Confocal imaging was performed on a Zeiss LSM 900 confocal microscope equipped with 405nm, 488nm, 561nm and 640nm laser lines using a Plan-Apochromat 20x/0.8 M27 objective and Airyscan 2 GaAsP-PMT detector. Identical laser intensity settings were applied to samples to be compared and equivalent Z-stacks were performed to capture the cell thickness. After acquisition, a maximum intensity projection of the Z-stack was applied using ZEN Blue 3.0 software. Nuclear colocalization between BCL6 and DAPI, and NFκB and DAPI, was analyzed in Imaris.

### Flow Cytometry: Endothelial Cell Activation

Confluent endothelial monolayers in a 24 or 48 well plate were pre-treated with inhibitors for 30min and stimulated with TNFα (20ng/mL final concentration) in M199+10% FBS for the indicated times. After treatment, conditioned medium was removed, cells were washed once with 1X PBS without Ca^2+^ or Mg^2+^, detached with room temperature Accutase, rescued with FACS buffer and pelleted at 360*xg* for 6min. Resuspended cells were stained with the flow cytometry antibody cocktail: ICAM-1-AF488 (Biolegend #32714), E-selectin-PE (Biolegend #322606), VCAM-1-APC (Biolegend #305810), BST2-PE/Cy7 (Biolegend #348416), and HLA-ABC-BV510 (Biolegend #311436) for 45min in the dark at 4⁰C, then washed with FACS buffer. Dilution information is available in the Supplemental Methods. ≥3,000 total events per sample were acquired on a BD Fortessa flow cytometer and data were analyzed in FlowJo. Debris were gated out by FSC/SSC and median fluorescence intensity of the live gate population was determined for each compensated fluorophore.

### Flow Cytometry: Leukocyte Adherence Assay

Peripheral blood mononuclear cells (PBMC) were enriched from the whole blood of healthy volunteers collected into ACD tubes, by Ficoll-Hypaque density centrifugation, or obtained from ATCC. After stimulation of endothelium, conditioned medium was removed and PBMC were added at an approximate ratio of 1EC:3PBMC, and allowed to adhere for 45min at 37⁰C. Nonadherent cells were removed with two washes with HBSS and one final wash with PBS without Ca^2+^ or Mg^2+^, followed by detachment of all remaining firmly adherent PBMC and endothelium with Accutase. Detached cocultures were pelleted and stained for flow cytometry: CD11a-FITC (BD #555383), CD105-BV510 (BD #747746) and CD3-APC/Fire 750 (Biolegend #317352). Antibody dilution information is available in the Supplemental Methods. 10,000 total events were acquired on a BD Fortessa flow cytometer and data were analyzed in FlowJo. Frequencies of PBMC in the adherent coculture fraction were determined by gating on CD105^neg^CD11a^+^ PBMC relative to the total live cell population.

### Fluorescence Leukocyte Adherence Assay

PBMC were labeled with CFSE at 5μM for 15min, then washed and resuspended in M19+10% FBS at #x10^6^ cells/mL. Confluent endothelial cells seeded in a clear bottom, black well 96 well plate and stimulated for the indicated times. After removal of stimulation medium, CFSE-labeled PBMC were added to endothelial cells and allowed to adhere for 45min at 37⁰C. Nonadherent cells were removed by decanting and washing twice with HBSS, then monolayers were fixed with 10% neutral buffered formaldehyde and GFP fluorescence and images were read on a Cytation5 microplate reader.

### HDAC Activity Assay

Primary endothelial cells were cultured to confluence in a 96 well plate, and treated with BCL6 inhibitors, TSA and/or TNFα for the indicated times. Histone deacetylase activity was measured using the In Situ Histone Deacetylase Activity Fluorometric Assay (Millipore Sigma) according to the vendor’s protocol. Fluorescence (ex/em) was read on a Cytation5 microplate imaging reader (Biotek). Blank well fluorescence was subtracted, and RFUs were normalized to a standard curve per vendor’s instructions.

### Analysis of Public Transcriptome Datasets

**Supplemental Table 1** provides a list of public datasets analyzed in this study.

### Statistical Analyses

## Results

### BCL6 is expressed in endothelial cells from both human and mouse

Endothelial-specific transcriptome analyses have revealed numerous transcription factor candidates regulating endothelial lineage specification and site specific heterogeneity. Focusing on heterogeneity of the inflammatory response, we recently reported that the DNA binding protein *BCL6* was among several immune-related genes that are differentially expressed across endothelium from the aorta, coronary artery, cardiac microvasculature, renal glomerulus and liver microvasculature [4].

Because BCL6 is primarily described in the lymphoid and myeloid compartments, we asked how BCL6 expression compares in the non-hematopoietic cell lineages and organs. We first analyzed public datasets of bulk organ-specific transcriptomes. In GTEx, BCL6 expression was broadly distributed, with the highest *BCL6* expression in whole blood and skeletal muscle (**Supplemental Figure 1**). *BCL6* expression was also widely detected across tissue types in developing human organs from Descartes [49]. We then asked how BCL6 expression compares in vascular endothelial cells with other cell lineages. In a public dataset of cultured human cells, GSE21212 ([50], n=2, GEO2R), *BCL6* expression was highest in B cells (5,541±1,500) and T cells (2,072±438), followed by smooth muscle cells (981±18); but was detectable in endothelial cells from all anatomic sites tested as well (**Figure 1a**). In GSE21212 ([50], n=2, GEO2R), *BCL6* was significantly differentially expression in large artery compared with umbilical vein endothelium, particularly large artery endothelium from coronary artery (714±22), pulmonary artery (652±40) and aorta (567±12) had higher expression levels than microvascular EC (303±45) or umbilical vein EC (347±14.5) (**Figure 1a**). In three other datasets (GSE131681 [51]; GSE3874 [52]; R2 RNA Atlas), among human endothelial cells from different tissues, endothelial cells from the heart and lung ranked most highly for *BCL6* expression (**Figure 1b, 1c, 1d**). For example, in the Human Endothelial Epigenome Database ([51], GSE131681, n=2-4), endocardial endothelium, aortic and pulmonary endothelial cells had the highest *BCL6* expression (p<0.0001) compared with coronary artery, carotid artery, renal artery, umbilical vein and saphenous vein endothelium (**Figure 1b**).

**Figure 1.**
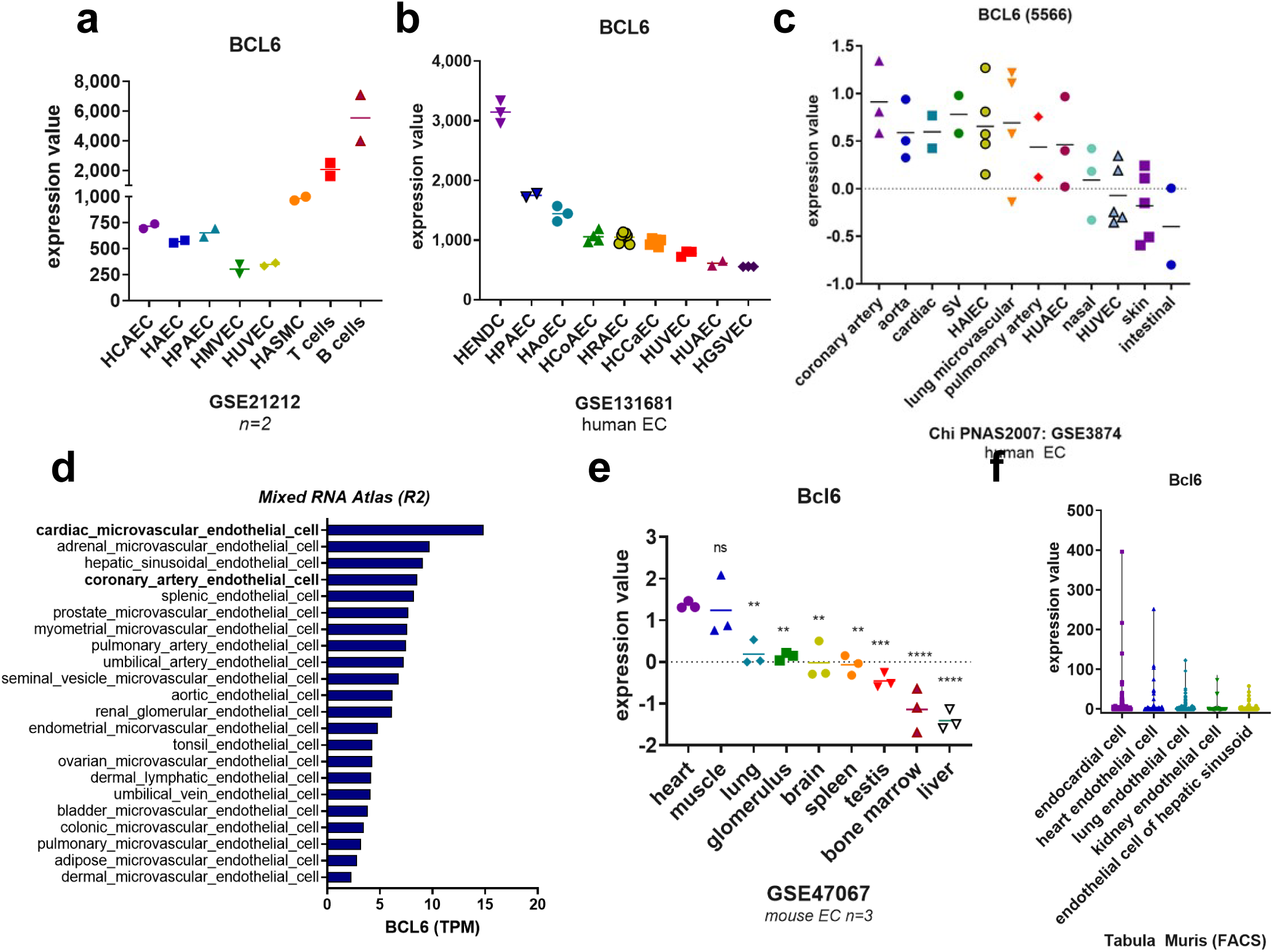
BCL6 is widely expressed, including in endothelial cells; and among endothelial cells is enriched in cardiac EC. *BCL6* expression is shown for cultured human endothelial cells from different anatomic sites compared with smooth muscle cells, T cells and B cells from GSE21212 (a) and across vascular beds from GSE131681 (b), GSE3874 (c), and the Mixed RNA Atlas (d). *Bcl6* expression in mouse endothelial cells isolated from different organs and analyzed in bulk from GSE47067 (e) and analyzed by single cell RNA-Sequencing in Tabula Muris (f). HAEC: human aortic endothelium; HCAEC: human coronary artery endothelium; HPAEC: human pulmonary artery endothelium; HMVEC: human microvascular endothelium; HRAEC: human renal artery endothelium; HCCAEC: human carotid artery endothelium; HUVEC: human umbilical vein endothelium; HUAEC: human umbilical artery endothelium; HGSVEC: human great saphenous vein endothelium; HASMC: human arterial smooth muscle cells

The results were confirmed in murine endothelial heterogeneity datasets. Among cultured mouse endothelium from 9 tissues, cardiac and muscle-derived endothelium had the highest *Bcl6* expression (**Figure 1e**, GSE47067 [53]). In mouse endothelium (Nolan, GSE47067, bulk EC from organs n=3), *Bcl6* was significantly higher in the heart compared with lung (p=0.0285), kidney glomerulus (p=0.0203), brain (p=0.0077), testis (p=0.0005), spleen (p=0.0057), bone marrow (p<0.0001) and liver (p<0.0001), but not muscle (p>0.999) [two way ANOVA, Tukey’s]. Like the heart, muscle endothelium also showed significantly higher levels than other tissues. Cleuren et al. used an EC-TRAP approach for the endothelial translatome ([54], GSE138630, n=3). Here, *Bcl6* was lowest in endothelium from the liver and brain; and higher in EC from lung and heart. Similarly, in mouse microvascular endothelium from heart, liver, and brain ([55]; GSE48209), *Bcl6* was significantly differentially expressed comparing heart and liver EC (p=0.001538, FC 0.5215, n=5 each).

Looking with higher resolution of the cell types within discrete organs, single cell sequencing of mouse tissues in Tabula Muris [56] showed that *Bcl6* was detectable in endocardial and endothelial cells of the heart, as well as smooth muscle cells and fibroblasts. In the lung, *Bcl6* was expressed in resident B cells, dendritic cells, stromal cells, type II pneumocytes and endothelium; in the kidney, *Bcl6* transcript was detected in all cells except collecting duct; and in the liver, *Bcl6* was highest in hepatocytes and Kupffer cells, but detectable in all cell types (**Figure 1f**). Lastly, comparing mouse endothelium in the single cell RNA dataset *Tabula Muris* [56], *Bcl6* was significantly higher in heart endocardial endothelium compared with other heart EC (p<0.0001). But, both cardiac endothelial cell types were significantly higher than EC from lung (p<0.0001), kidney (p=0.0383) or hepatic sinusoid EC (p<0.0001) (**Figure 1f**).

Similarly, in the human *Tabula Sapiens* database [57], BCL6 expression was strongest in a subset of the immune compartment, but was also widely detected throughout other cell lineages including endothelium (**Figure 2a**). At the single cell level, in *Tabula Sapiens*, human endothelial cells showed a distribution of expression, with a subset of highly expressing endothelium that were only categorized as of “vasculature” origin (**Figure 2b**). We segmented the endothelial cells based on *BCL6* expression ≥1 and <1, then compared top differentially expressed genes in the two groups. *BCL6*+ expression was correlated with expression of *NFKBIA*, *NFKBIZ*, *TNFAIP3* (negative NFΚB regulators), *ICAM1* (adhesion molecule), *JUNB* (component of AP-1 transcription factor complex), and *MAFF* (an endothelial master regulator TF). In contrast, the BCL6^lo^ endothelial cells had higher levels of *CAVIN2, ID1* (a transcriptional repressor), *IFITM3* (IFN/anti-viral signaling), and *TXNIP* (a regulator of VEGFR2 signaling) (**Figure 2c**).

**Figure 2.**
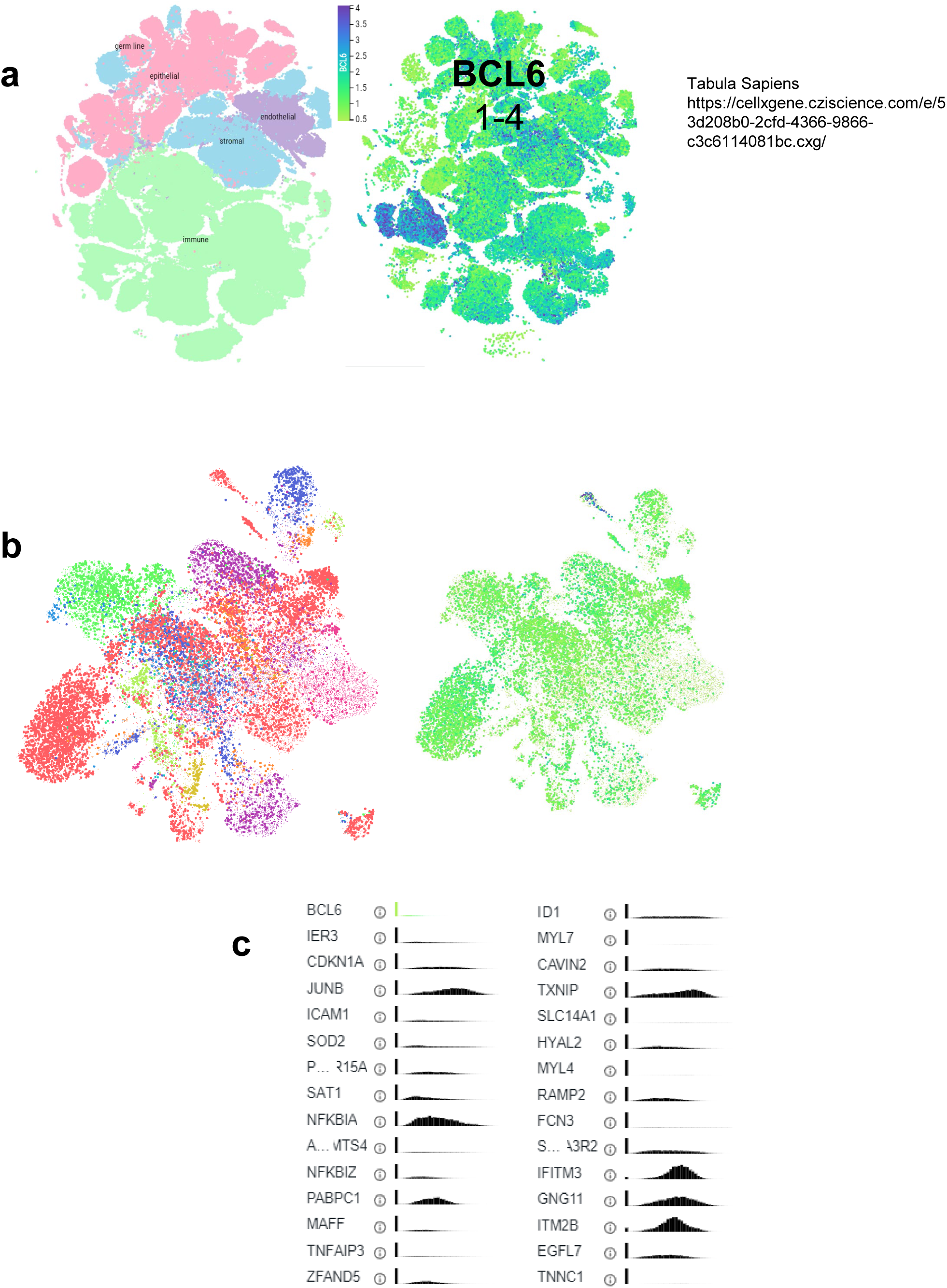
Expression of BCL6 across human cell types from the single cell gene expression atlas Tabula Sapiens, across cell types and organs (a), and within endothelial cells from different organs and tissues (b). Top marker genes and expression distribution in endothelial cells segmented by positive BCL6 expression (left, 1+) and negative BCL6 expression (right, <1).

Therefore, BCL6 is expressed across endothelial cells from many different anatomic sites. Moreover, our analyses demonstrate that basal BCL6 expression in endothelium was organotypic, with particularly enriched expression in cardiac endothelium.

#### The chromatin landscape around BCL6 is highly accessible in endothelium

Because BCL6 was lowly but basally expressed in endothelium, we examined the histone marks around the BCL6 gene. Super-enhancers characterized by high H3K27Ac signal are important for and localize around genes controlling cell identity [50, 56, 58–61] [62, 63]. We analyzed the chromatin architecture in public datasets from human endothelial cells [[64, 65] [66] [51]], and verified that *BCL6* gene exhibits high chromatin accessibility, with a chromatin landscape indicative of active/poised enhancers (**Supplemental Figure 2a**). Concordantly, in a mouse dataset [67], the *Bcl6* gene showed high chromatin accessibility by ATAC-Seq in endothelial cells from the lung, liver and kidney (**Supplemental Figure 2b**).

To confirm endothelial BCL6 expression at the protein level, we stained normal human FFPE tissue for BCL6 and CD31 by immunofluorescence (n=4 donors). Positive control tissue human tonsil stained reliably in germinal centers as expected, confirming antibody specificity (**Figure 3a**). BCL6 was detected in normal human kidney, but mostly as diffuse staining in tubules and only rarely in vascular compartments (glomeruli or peritubular capillaries). In human lung, occasional BCL6 positive nuclei were observed, within CD31+ endothelial cells. In normal human heart and liver tissue, however, there was scant BCL6 staining (**Figure 3b**). These results were confirmed in the Human Protein Atlas, which showed nearly identical staining patterns using two different antibody clones (**Supplemental Figure 3**). Therefore, BCL6 was expressed at the mRNA level but not consistently detected at the protein level in cardiac endothelium *in situ*.

**Figure 3.**
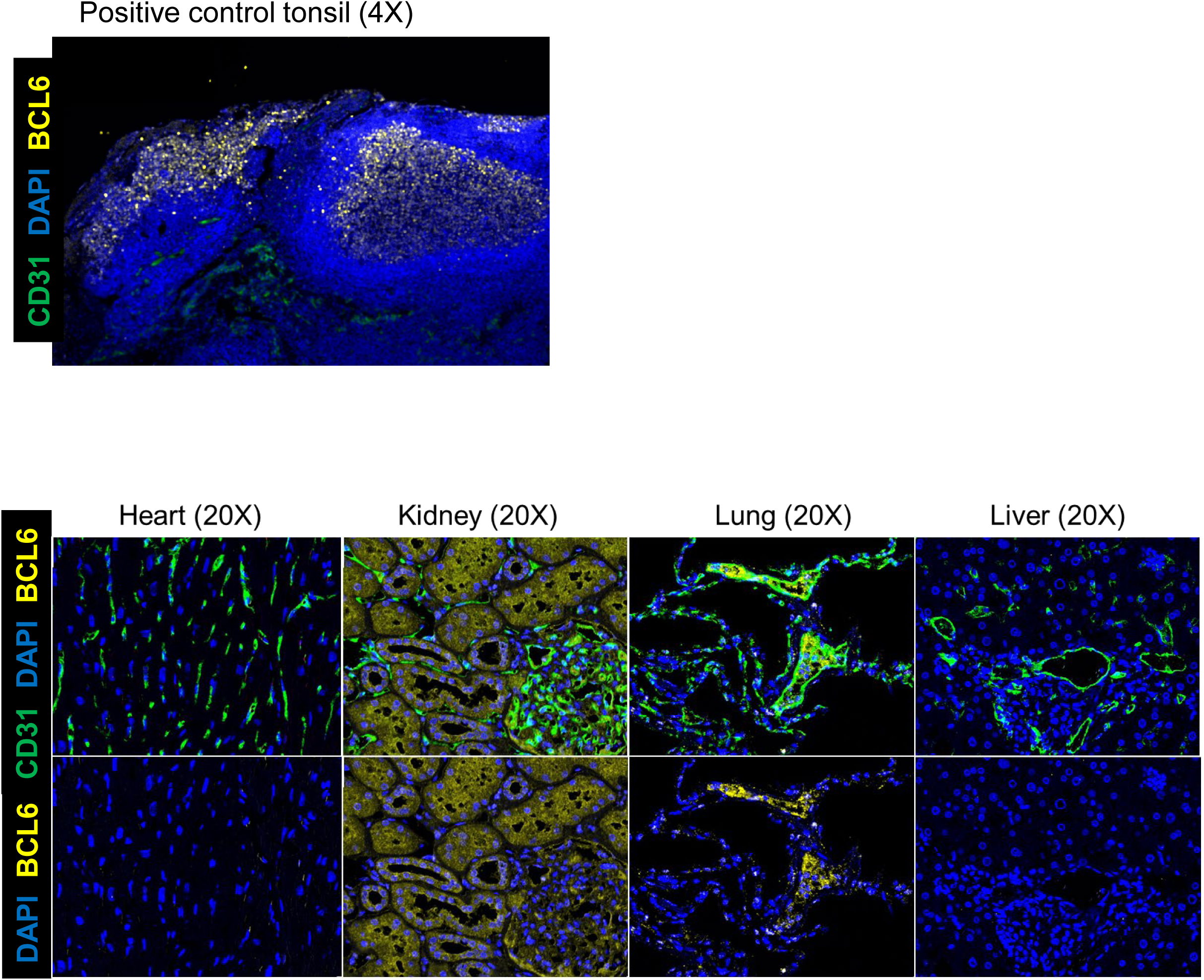
BCL6 protein expression in normal human heart, lung, liver and kidney. FFPE remnant tissue from normal human organs from 5 different donors was probed by multiplex immunofluorescence for CD31 and BCL6. Representative micrographs show staining.

Thus, the BCL6 gene is demonstrates low basal mRNA and protein expression but high chromatin accessibility in endothelium. That the *BCL6* gene is downstream of a superenhancer in EC, suggests it may be poised as an immediate early response gene in endothelial inflammation.

### BCL6 is upregulated under TNFα stimulation in endothelium

It has already been reported that BCL6 is upregulated in EC by angiogenic stimuli [30], but studies in immune cells have shown that several cytokines also upregulate BCL6. We next asked if BCL6 was cytokine inducible in endothelial cells. To determine under what other conditions BCL6 expression is increased in endothelium, we examined BCL6 transcript variation across endothelial transcriptome datasets (EndoDB) and by querying GEO profiles for conditions with differential expression of BCL6 in endothelium (**Figure 4a**). Myriad stimuli altered BCL6 expression in EC. Of particular interest, endothelial BCL6 expression was significantly increased in 3 unique datasets of TNFα stimulation (GSE6257, GSE2639, GSE144803) (**Figure 4a**).

**Figure 4.**
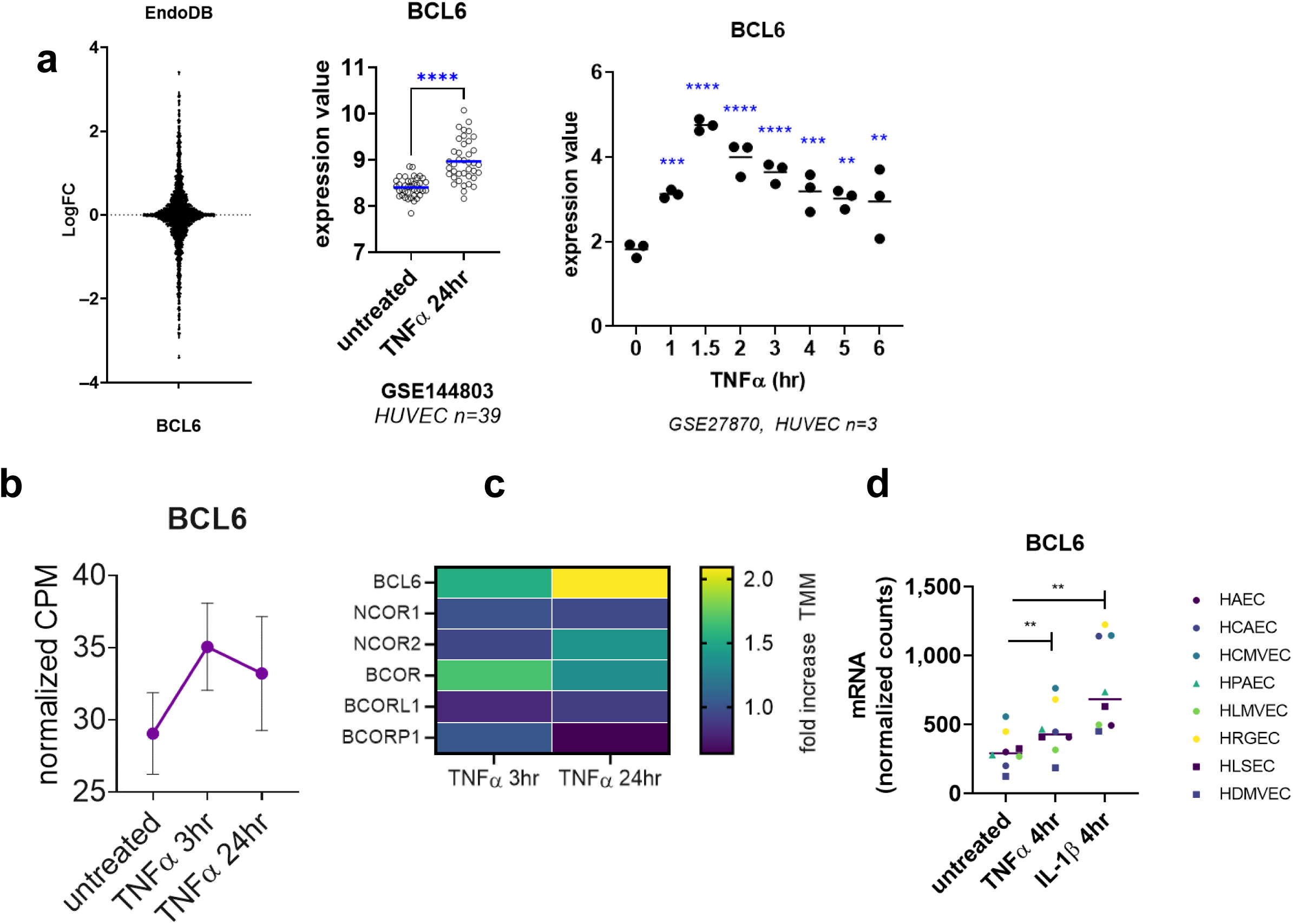
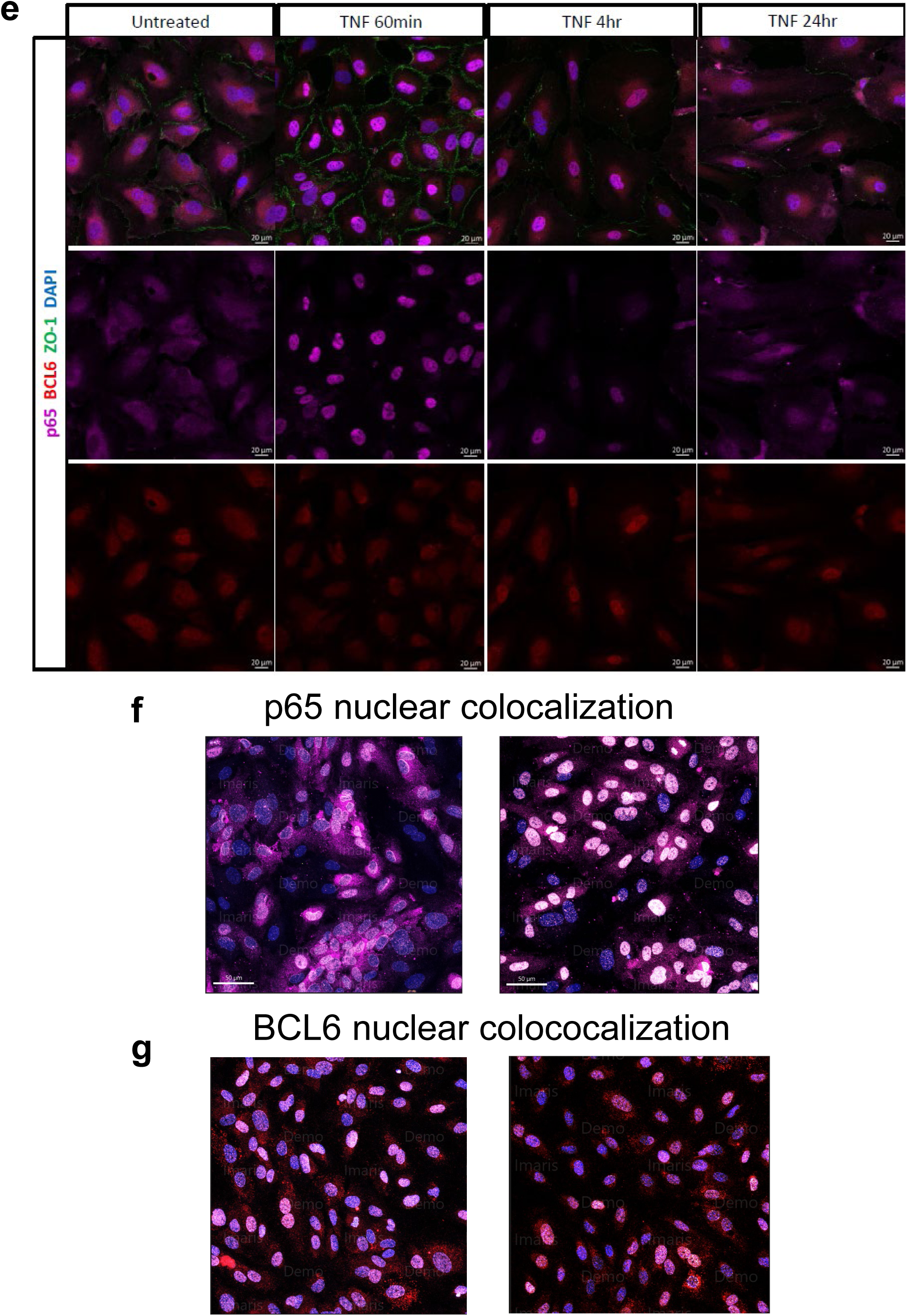
BCL6 expression is inducible by TNFα in endothelium. (a) Left panel: Variation in BCL6 expression across expression datasets in EndoDB. Middle panel: BCL6 expression after 24hr of TNFα stimulation across 39 different HUVEC donors, queried in GSE144803 from GEO. Right panel: Expression data from GSE27870 in HUVEC (n=3 replicates) was queried from GEO. (b) Expression of BCL6 over time measured by RNA-Seq in primary HAEC (n=3-4 biological replicates). (c) Fold increase in transcript abundance of BCL6 and related corepressor genes is shown in the heat map. (d) Expression level of BCL6 at 4hr across multiple different endothelial cell types (measured by Nanostring). BCL6 intracellular localization in endothelium is predominantly nuclear, and does not change under TNFα stimulation. Confluent HAEC were treated with TNFα for the indicated times, and then stained for intracellular p65 NFκB and BCL6. (e) shows exemplary micrographs. (f) and (g) show intensity of signal colocalization between DAPI and p65, and DAPI and BCL6 in untreated (left panels) and TNFα-treated (right panels) HAEC.

To confirm this, we treated primary human aortic endothelial cells (HAEC n=3 biological replicates) and primary human endothelium from 8 different vascular beds with TNFα, and measured BCL6 expression. In HMEC-1 cells, BCL6 expression was upregulated by 3.95-fold within 2-3hr of stimulation with TNFα (**Supplemental Figure 4**) and maintained at 2.18-fold to 24hr. In HMEC-1, BCL6 tempo of induction paralleled transcript expression of known NFκB responsive genes like *TNFAIP3* and *NFKB1* (**Figure 4b**). This was confirmed in HAEC from three different donors (**Figure 4c**), and BCL6 was upregulated across endothelial cells from distinct vascular beds (**Figure 4d**). Our *in vitro* results therefore corroborate public transcriptome datasets that TNFα stimulated endothelial cells upregulate BCL6 expression.

Because BCL6 has been described as a transcriptional repressor, we examined localization of BCL6 and NFκB p65 protein in untreated and TNFα stimulated endothelium by immunofluorescence. As expected, NFκB was predominantly cytosolic in untreated EC, but clearly moved to the nucleus within 30min of TNFα stimulation (**Figure 4e, 4f**). NFκB nuclear localization was decreased by 3hr post stimulation, and by 24hr was predominantly cytoplasmic (**Figure 4e**). In contrast, BCL6 protein was detected as nuclear staining in untreated endothelium, with some perinuclear cytosolic staining also observed (**Figure 4e, 4g**). Pre-formed BCL6 nuclear localization was maintained at 15min and 30min of TNFα stimulation (**Figure 4g**). Because BCL6 was upregulated after TNFα stimulation, we tested BCL6 localization after 4hr and 24hr of activation. Nuclear BCL6 signal was maintained at 4hr and 24hr (**Figure 4e**).

Therefore, BCL6 expression is rapidly increased in endothelial cells by TNFα treatment, through NFκB, but unlike NFκB, remains predominantly nuclear over the course of TNFα stimulation.

### BCL6 upregulation by TNFα is NFκB-dependent

We asked what transcription factors might regulate BCL6 expression in endothelial cells. To that end, we queried databases of transcription factor binding to the BCL6 gene (MotifMap, CHEA, TRANSFAC, ENCODE, and CistromeDB). Interestingly, RELA (NFκB) was predicted or confirmed to bind the BCL6 gene by 3 different approaches, including a query filtered on empiric transcription factor ChIP-Seq data in endothelium (**Figure 5a**).

**Figure 5.**
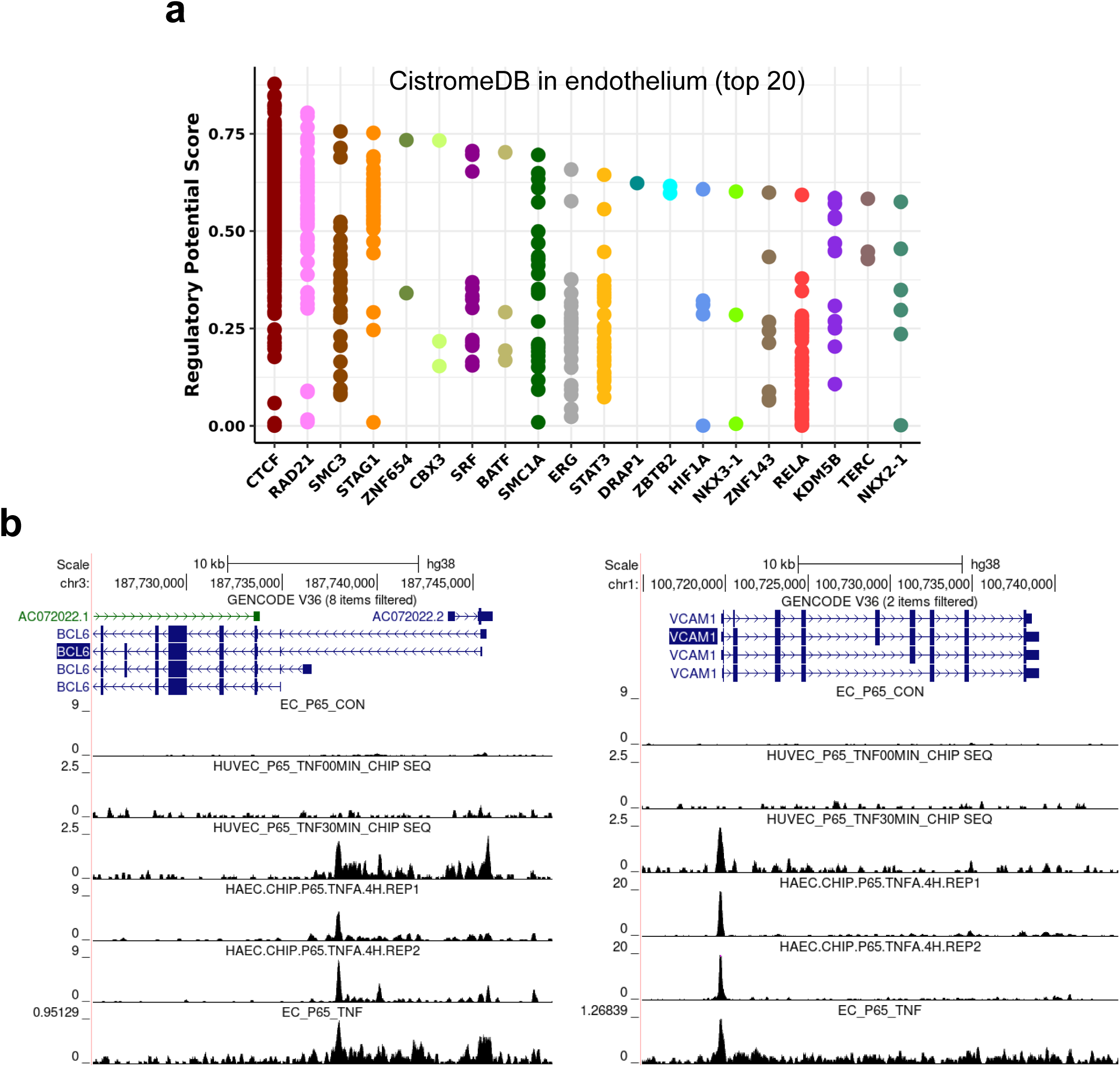
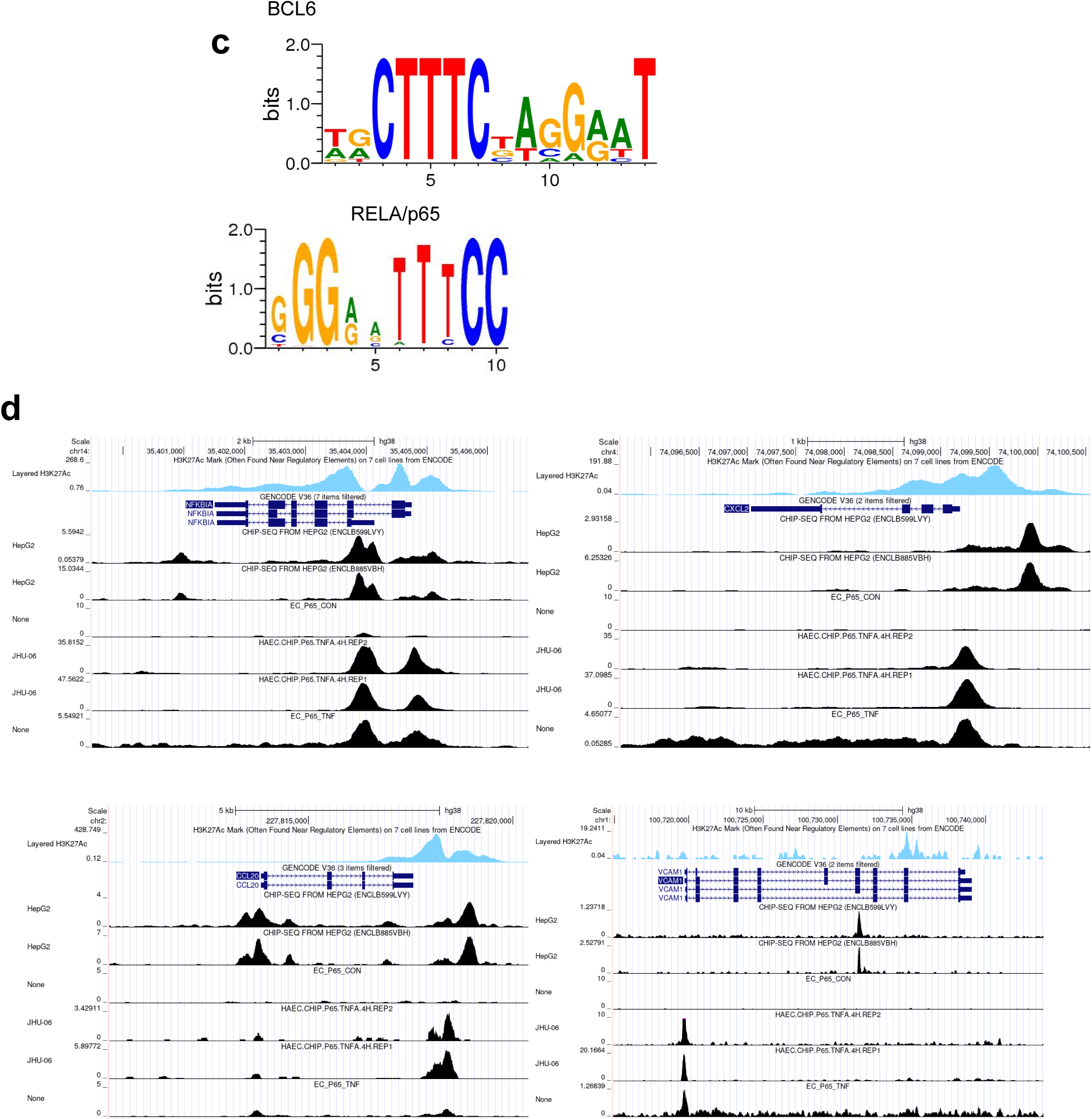
Transcription factors and chromatin modifiers binding within +/-10kb of the TSS of the BCL6 gene were queried in Cistrome DB (a). ChIP-Seq tracks of p65 NFKB binding from (Hogan), whosing that TNFα treatment of EC increases p65 NFkB binding at the *VCAM1* and *BCL6* genes (b). BCL6 binds to NFκB target genes. (c) BCL6 consensus DNA binding motif partially overlaps with RelA. (d) Genome browser views of H3K27Ac ChIP-Seq in HUVEC (ENCODE), BCL6 ChIP-Seq tracks from HepG2 cells (ENCODE), and p65 NFKB ChIP-Seq in unstimulated and TNFα-treated HAEC (Hogan).

Examining NFκB p65 ChIP-Seq data in HAEC and HUVEC under TNFα stimulation from three different studies [66, 68, 69], we identified that NFκB does not bind to BCL6 gene in unstimulated endothelium. However, RELA binding is highly increased in endothelial cells treated with TNFα or IL-1β [66]. For example, minimal p65/RelA binding to the BCL6 gene was detected in untreated EC, but increased more than 8-fold under TNFα stimulation (**Figure 5a**). Accordingly, the BCL6 gene contains putative NFκB and REL transcription factor binding sequences (HOMER). In contrast, no peaks for other transcription factors like IRF1 or Jun were detected around the *BCL6* gene in either untreated or TNFα stimulated EC (*data not shown*).

### Transcription factor similarity with NFκB

Since BCL6 is a DNA binding protein, we analyzed its putative gene targets. Using GOMo (Gene Ontology for Motifs), which scans gene promoters for the queried DNA motif and functionally annotates the clusters, we analyzed the GO terms significantly associated with genes associated with the human MA0463.2 and mouse MA0463.1 BCL6 motifs [70], to find a predominance of genes categorized as inflammatory response genes (GO:0006954). Further, matrix alignment of the BCL6 and Rel/NFκB DNA binding motifs showed partial overlap between consensus sequences in a tiled manner (**Figure 5c**).

To address the possibility that BCL6 regulates NFκB-dependent genes in endothelium, we examined the predicted and defined target genes bound by BCL6 for genes that are critical for endothelial inflammatory responses. Using DNA motif analysis, MotifMap and Transfac, we found that BCL6 was predicted to bind to numerous adhesion molecule and chemokine genes that are upregulated in endothelium by TNFα, including *VCAM1*, *CXCL2*, *CX3CL1*, *SELE*, and *CCL5*; as well as NFKB genes themselves (**Table 1**, exemplary peaks in **Figure 5d**). We also explored the ChIP-Atlas (https://chip-atlas.org/target_genes) for genes with empirically defined BCL6 binding +/-5kb across available datasets (B cells). Among the highest scoring hits were *SOCS2*, *MAPKAPK3* and *CISH*, as well as *BCL6* itself; but also *CXCL2*, *CXCL11*, *CCL5*, *CD274*, *NFKBIA*, *NFKBIB*, *NFKBIZ*, and *TNFAIP3*. Additionally, we queried gene targets of BCL6 specifically for endothelial inflammatory response genes, and found significant peaks at *CCL20*, *VCAM1*, *CX3CL1*, and *CXCL1*.

**Table 1.** Genes with proximal binding of BCL6, from ChIP-Seq data (ENCODE) in HepG2 cells.

| At TSS | Intragenic | Other<br>(up/down<br>stream) |
| --- | --- | --- |
| ICAM1 | VCAM1 | CXCL2 (up) |
| SOCS2 | GATA4 | CCL20 (dwn) |
| CCL20 | GATA6 | CSF1 (up) |
| TNFAIP3 | TNFAIP3 | CXCL5 (up) |
| HES1 | HES1 | RELA (dwn) |
| CX3CL1 | TNFSF4 |  |
| TNFSF4 | NFKBIA |  |
| NFKBIA | NFKBIZ |  |
| NFKB2 | NFKB2 |  |

To confirm the potential for BCL6 to bind to NFκB-regulated genes, we compared BCL6 genomic binding patterns [64] in HepG2, overlaid with endothelial chromatin accessibility (HUVEC H3K27Ac), to p65 NFκB binding in TNFα-activated endothelium [66] BCL6 binding sites indeed appeared to be slightly upstream or directly overlaid on many NFκB binding sites (**Figure 5d**). We observed several patterns: 1) BCL6 and RELA peaks partially or wholly overlapped at the same gene (*NFKBIA*, *TNFAIP3*, *CX3CL1*, *CCL20*, *ICAM1*, *NFKB2*, *STAT1*, *IL32*, *SOCS3, CD274*); 2) BCL6 and RELA both bound a gene but at different locations (*VCAM1*, *SOCS2*, *CXCL2*, *CLU*); or 3) genes were bound only by RELA (*SELE*, *CCL5*, *CXCL10*, *CCL2*, *CD40*, *MIR155HG*, *CXCL8*, *NFKB1*, *CSF1*, *CSF2*, *IKBKE*) but not BCL6. Exemplary genes *CXCL2*, *CCL20, VCAM,* and *NFKBIA* are shown (**Figure 5d**).

These analyses demonstrate that BCL6 and p65 NFκB putatively bind to shared target genes involved in the endothelial cell response to TNFα.

### Function of endothelial BCL6 in NFκB-driven inflammation

To understand the functional effects of BCL6 in endothelium, we established stable knockdown and overexpressing cell lines using lentiviral transduction of HAEC with shRNA or the open reading frame of BCL6. TeloHAEC consistently expressed the GFP reporter (**Supplemental Figure 5a**), and BCL6 overexpression was readily detected at increasing MOI in primary HAEC as well (**Supplemental Figure 5b**). Passaged transduced TeloHAEC also showed expected overexpression and reduced expression of *BCL6* (**Supplemental Figure 5c**). The immortalized aortic endothelial cell line TeloHAEC reliably responded to TNFα by upregulating adhesion molecules (**Supplemental Figure 5d**), as reported [71] demonstrating its utility in this model.

To understand the global transcriptional alterations by BCL6, we performed RNA-Seq on immortalized TeloHAEC transduced with GFP or BCL6 open reading frame for overexpression, or with GAPDH or BCL6 shRNA for knockdown; and confirmed in primary HAEC treated with a BCL6 degrading inhibitor BI-3802 [38].To confirm specificity for BCL6, we first examined the effects of BCL6 depletion on previously described BCL6 gene targets [72, 73], such as *SOCS2*, *LDLR*, and *FABP4* [74]. BCL6 overexpression dose-dependently suppressed *SOCS2* and *FABP4*. In total 182 genes were downregulated by overexpression of BCL6, including *BMPER*, *F3, CXCL8,* and *SELP* (**Supplemental Figure 5e**). Conversely, loss of BCL6, via the degrading inhibitor BI-3802 or shRNA, augmented *SOCS2* and *LDLR* expression, with or without TNFα treatment (**Supplemental Figure 5f** and *data not shown*). Similar results were obtained when primary HAEC were transfected with siRNA against BCL6, although only a 50% reduction in BCL6 was achieved with siRNA (**Supplemental Figure 6**).

In particular, overexpression of BCL6 dose-dependently suppressed basal expression of *IL6, CXCL8*, *CXCL1*, *SELP,* and *SELE,* suggesting endogenous repression of these genes in unstimulated cells. On the other hand, 51 genes were highly upregulated (CPM>2.0, >5-fold) by BCL6 shRNA in untreated cells, while 106 genes were downregulated (<50%). Of interest, knockdown of BCL6 alone significantly increased chemokines *CXCL1*, *CXCL2*, *CSF2*, *CSF3*, *CCL2*, *CCL5, CXCL10* and *CXCL11*, and adhesion molecules *BST2*, *SELE,* and *ICAM1*. Concordantly, secretion of chemokines RANTES (*CCL5*) and IP-10 (*CXCL10*) was increased by transduced, unstimulated ECs (RANTES: 2-fold, IP-10: 4-fold, *not shown*), as was cell surface BST2 expression. Further, BCL6 shRNA augmented, while overexpression reduced, a number of basal genes, including *CCL2*, *INHBA, BST2*, *CXCL1*, and *SELE*.

When transduced cells were treated with TNFα, BCL6 overexpression reduced TNFα-induced chemokines *CXCL10*, *CXCL11*, and *CX3CL1*, while their gene expression was conversely augmented by BCL6 shRNA (**Figure 6a, 6b**). The adhesion molecule *BST2* was similarly increased by BCL6 knockdown (**Figure 6a**). Interestingly, *VCAM1* mRNA was decreased by BCL6 shRNA (**Figure 6a**). In total, 51 TNFα-induced genes (CPM>2.0, >2-fold) were reduced (<60%) by BCL6 overexpression, including: *TNFSF10*, *TNFSF15*, *TNFSF9*, *IFIT1*, *IFIT2*, *IFITM1*, *ISG15, CX3CL1, CXCL10, CXCL5,* and *BATF2*. Reciprocally, 79 TNFα-induced genes (>2-fold) were further increased (>50%) by BCL6 shRNA, including: *CXCL11, CSF3, BST2, BATF3, IFIT1, IFIT2, IFITM1,* and *TAP2*. Results for VCAM-1, E-selectin, and ICAM-1 were verified at the protein level by flow cytometry (**Figure 6c**).

**Figure 6.**
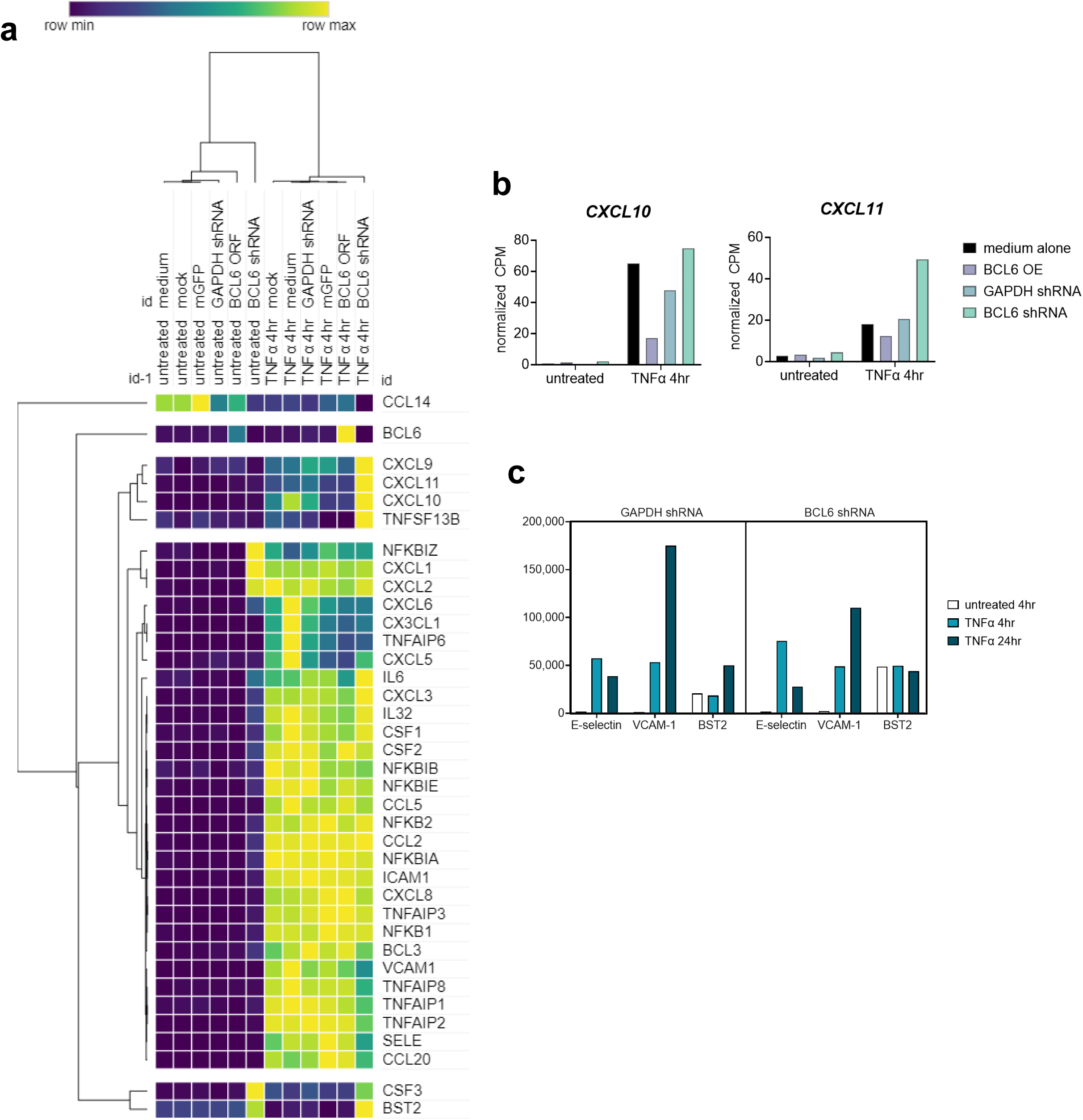
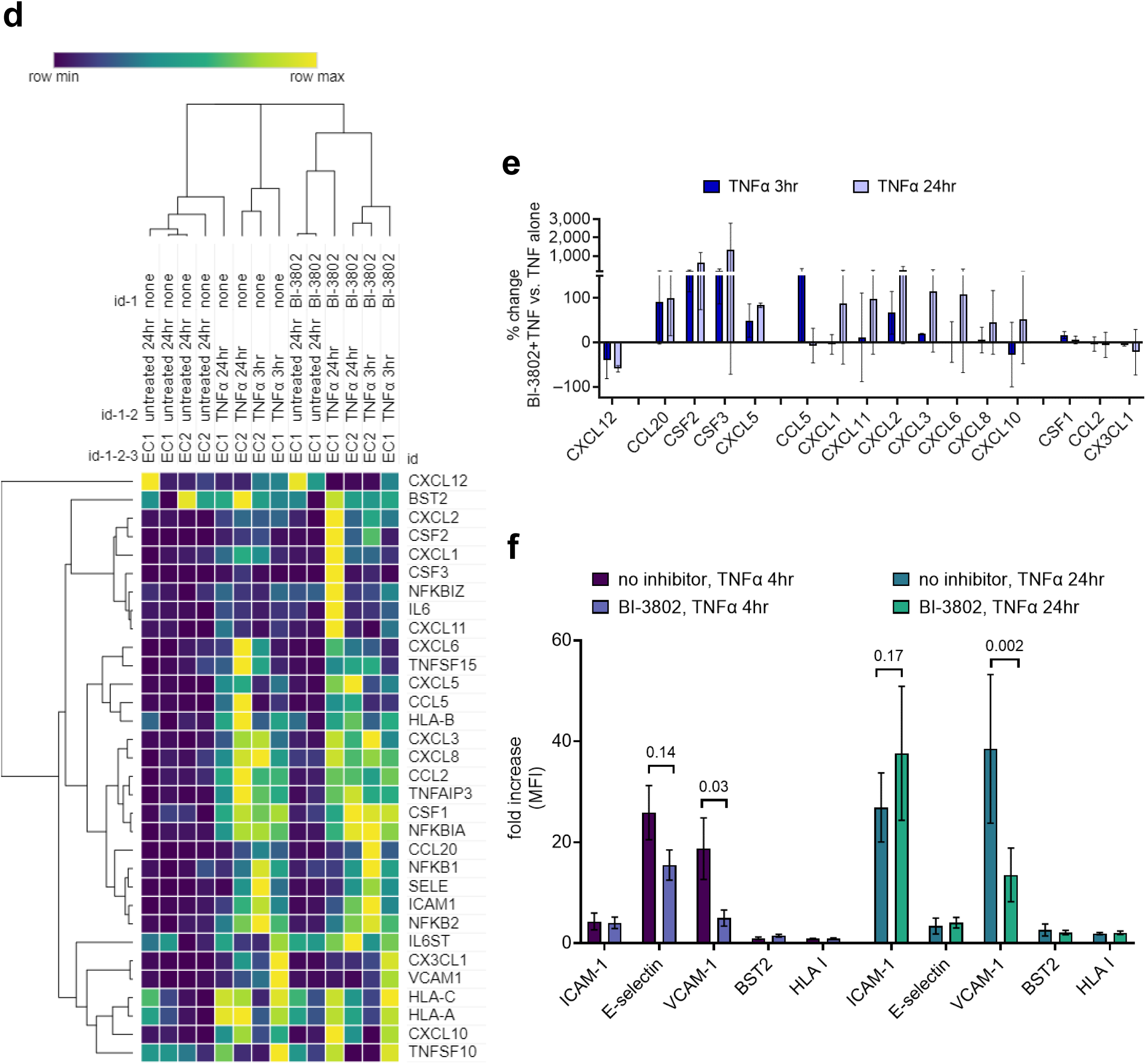
TeloHAEC were mock transduced or stably transduced with lentiviral vectors expressing GFP alone, BCL6 open reading frame, GAPDH shRNA, or BCL6 shRNA. Transduced and control cells were treated with TNFα for 4hr, then global mRNA expression was tested by RNA-Seq. (a) Heat map shows relative expression of TNFα induced genes. (b) Graphs show normalized CPM for transcript counts of representative genes. (c) Transduced endothelial cells were tested for cell surface adhesion molecule expression by flow cytometry. Primary human aortic endothelial cells (n=3 biological replicates) were pre-treated with the BCL6 degrader BI-3802, then stimulated with TNFα. Gene expression was measured by RNA-Seq. (d) Heat map shows relative gene expression. (e) Graph shows percent change in transcript abundance comparing BI-3802 treated HAEC to untreated HAEC, after TNFα stimulation. (f) Primary cardiac endothelial cells were tested for cell surface protein expression by flow cytometry. Results are graphed as fold increase in MFI for each marker (n=4-6 biological replicates). Group differences were analyzed by 2-way ANOVA followed by uncorrected Fisher’s LSD.

We then confirmed these results in primary human aortic endothelial cells pre-treated with the BCL6 degrading inhibitor BI-3802, for 4hr or 24hr (**Figure 6d**). As above, transcriptome changes were measured by RNA-Seq, and then tested by protein expression using flow cytometry (cell surface adhesion molecules) and Luminex/ELISA (secreted chemokines). The well-described TNFα-induced endothelial chemokines IL8/*CXCL8*, *CCL2*/MCP-1, and *CCL5*/RANTES were not affected by BCL6 depletion, nor were *ICAM1* or HLA class I genes (*HLA-A, B, C*) (**Figure 6d, 6e**). We have previously reported that *CXCL12* is downregulated by TNFα [75]; and this was not impacted by BI-3802. Other NFκB-inducible genes (*TNFAIP3*, *TNFAIP2*, *IKBKE*, *NFKB1/2*, *NFKBIZ*, *BCL3*, *IRF1*) were not changed in TNFα+BI-3802 conditions. Some genes were not affected by TNFα or BI-3802 alone, but were changed in the combination condition, including *C3* which was highly upregulated. However, other chemokines including *CXCL1*, *CXCL2* (MIP2α), *CSF2, CSF3, CXCL10*, *CXCL11*, *CXCL5*, and *CXCL6* were enhanced by BI-3802 (**Figure 6e**), as were cytokines *IL1A*, *IL6, IL15*. TNFα-induced expression of *SELE*/E-selectin was not affected at 4hr but was increased at 24hr. As with shRNA, *VCAM1* was reduced in the presence of TNF+BI-3802 compared with TNFα alone (**Figure 6d**). The effects of BI-3802 on TNFα-induced adhesion molecules was confirmed by flow cytometry, which showed that BCL6 degradation reduced E-selectin and VCAM-1 at 4hr, and reduced VCAM-1 at 24hr (**Figure 6f**).

These overexpression and depletion experiments therefore demonstrate an endogenous effect of BCL6 on NFκB-target genes in endothelial cells.

### BTB domain inhibitors suppress the NFκB-driven response to TNFα in endothelium

BCL6 suppresses gene expression in macrophages and B cells, in part via recruitment of HDACs via BCOR/NCOR corepressors that bind to its BTB domain. Several small molecule inhibitors and peptidomimetics have been developed that selectively target the BTB domain to disrupt complex formation with corepressors [37, 41]. Notably, these inhibitors do not dissociate BCL6 homodimers or inhibit BCL6 binding to target DNA. Since knockdown or degradation of BCL6 de-repressed a subset of NFκB response genes, we questioned whether inhibition of the BTB domain of BCL6 could potentiate the endothelial response to TNFα. This would implicate the corepressors NCOR/SMRT and/or BCOR, and histone deacetylases (HDACs) in BCL6-mediated repression of gene expression.

To answer this, we pretreated HAECs with three BTB inhibitors: low potency 79-6, its high potency derivative FX1[41], and a different molecule BI-3812 [37] prior to stimulation with TNFα (4hr, 24hr). First, we measured the effects of the BCL6 BTB inhibitor FX1 (100μM) on TNFα-induced mRNA by RNA-Seq (n=2-3 HAEC). Then we confirmed mRNA changes with Nanostring and protein changes by flow cytometry, Luminex and ELISA. Antagonism of the BCL6 BTB domain by FX1 or BI-3812 led to dramatic alterations in the TNFα-induced transcriptome in EC, while the FX1 analog 79-6 at an even higher dose had little effect (**Figure 7a, Supplemental Figure 7a, 7b**).

**Figure 7.**
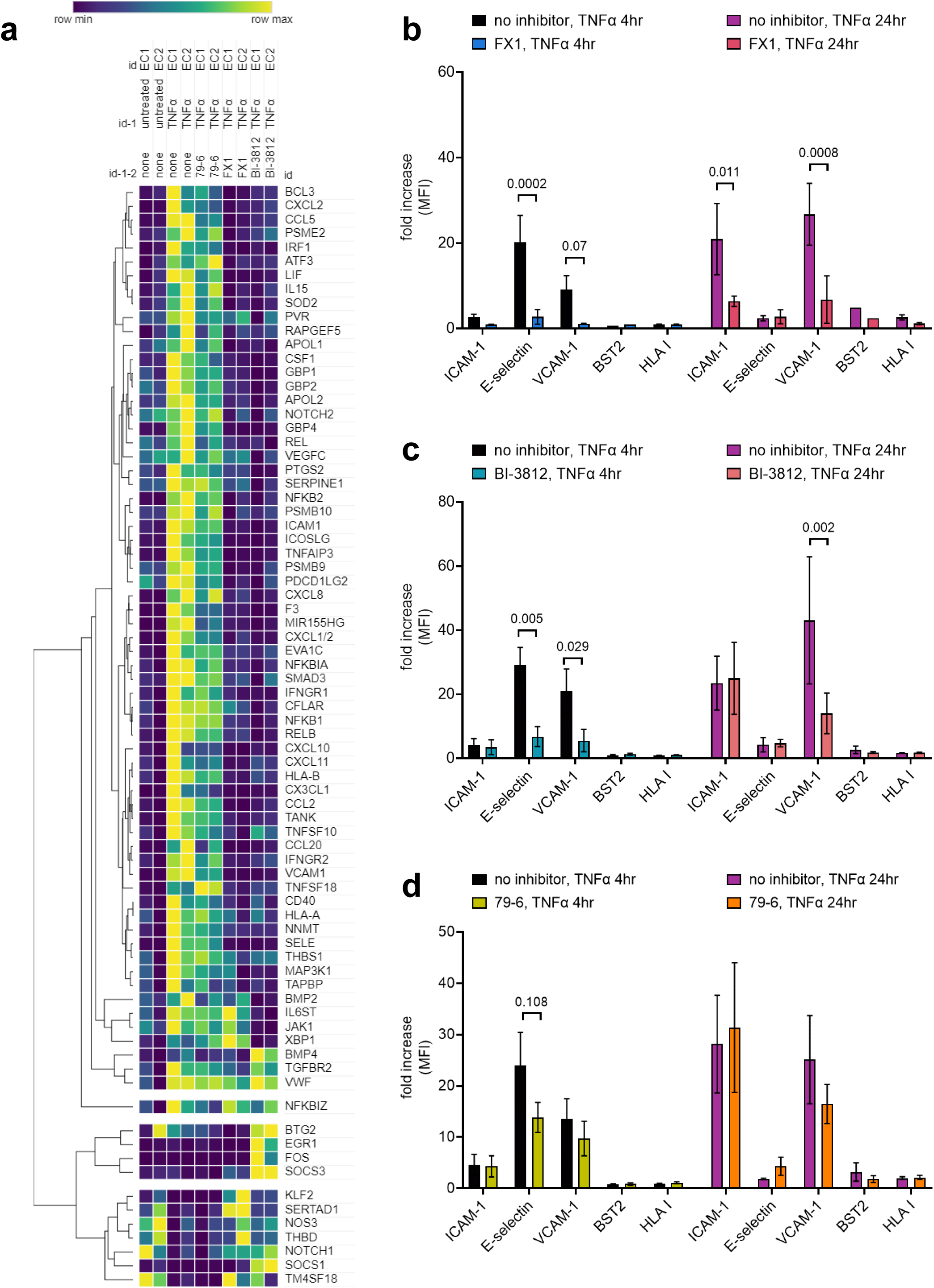
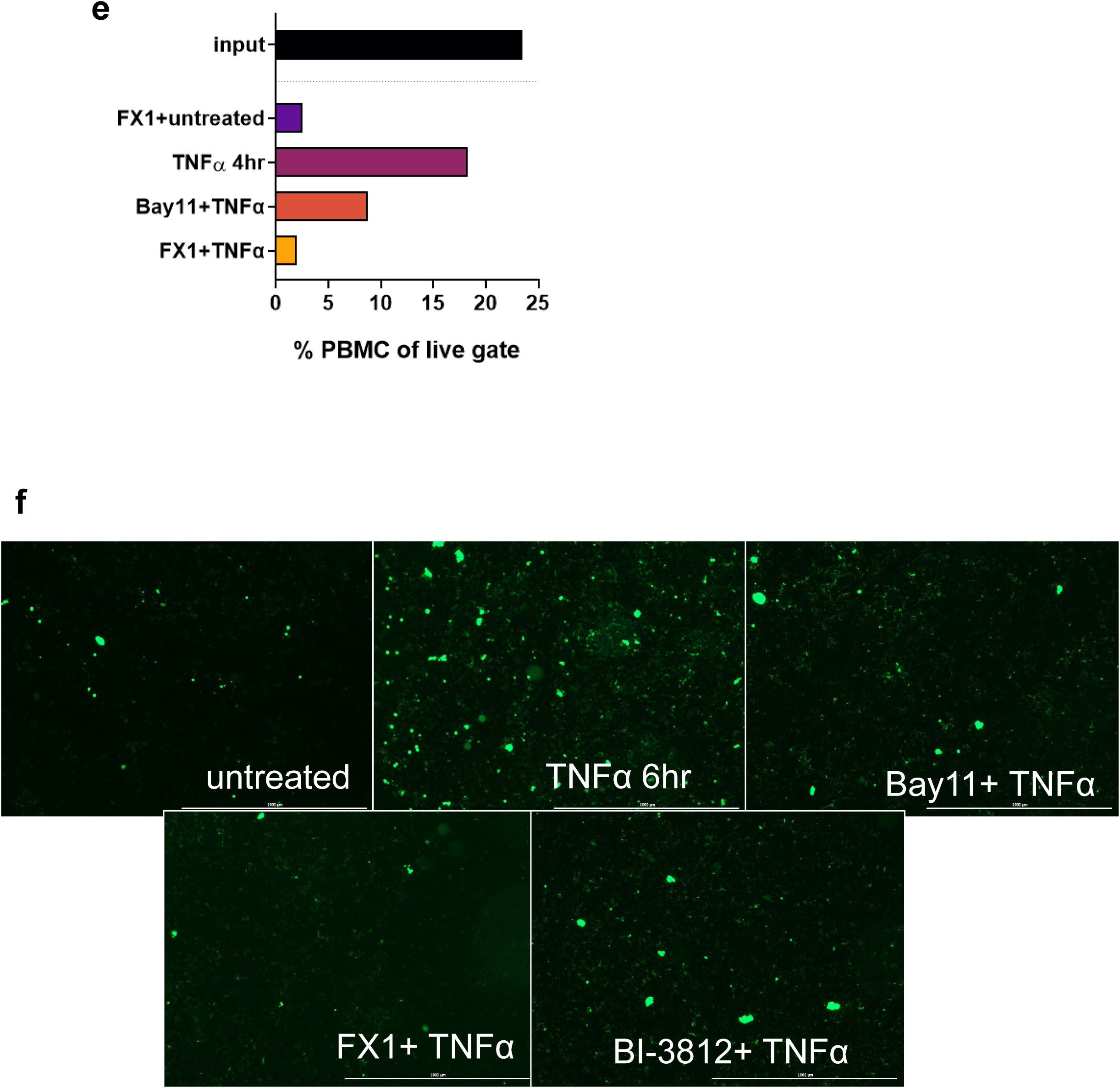
BTB domain inhibition affects endothelial pro-adhesive gene and protein expression. (a) Heat map from Nanostring showing 79-6, FX1 and BI-3812 (HAEC n=2 biological replicates). (b, c, d) Cell surface adhesion molecules were measured by flow cytometry on primary HAEC (n=3-6 biological replicates). BTB domain inhibition affects endothelial pro-adhesive gene and protein expression. (e) Conditioned supernatants from gene expression experiments were tested for secretion of chemokines and cytokines by multiplex Luminex assay. Groups were compared by two way ANOVA, followed by Dunnet’s multiple comparisons test. # p<0.1, * p<0.05, ** p<0.01, *** p<0.001 (n=3 biological replicates). BTB domain inhibition affects endothelial pro-adhesive gene and protein expression. (e) Allogeneic PBMC were cultured with to confluent conditioned endothelial monolayers. After washing, adherent cells were detached and the proportion of leukocytes among the adherent fraction was determined by flow cytometry. (f) CFSE-labeled U937 monocytes were cultured with conditioned endothelial monolayers. After washing, monocyte adherence was determined by microscopy.

As anticipated, some genes were similarly affected by the BCL6-BTB inhibitor as by BCL6 depletion, suggesting that regulation of those genes is BTB-dependent. For example, FX1 upregulated *CCL5* and *CXCL10* after TNFα activation. Conversely, some genes that were downregulated by TNFα treatment, such as *NOTCH1* and *EGR1*, were restored by the presence of FX1. And, BTB inhibitors had no effect on a subset of TNFα-induced transcripts such as *VWF, TGFBR2*, or *NFKBIZ*. 79-6, a lower potency BTB inhibitor, demonstrated less of an effect compared with FX1 and BI-3812.

Surprisingly, however, the presence of BTB domain inhibitors FX1 or BI-3812 significantly blocked TNFα-induction of many other genes, including *VCAM1* (**Fig 7a, 7b, 7c**) and *CXCL1/2* (**Fig 7a**). The effects of FX1 on TNFα-induced gene expression were confirmed with Nanostring, where FX1 suppressed more than half of TNFα-induced genes, including *CCL2*, *CCL5*, *CCL20*, *CXCL1*, *VCAM1*, *CXCL10*, *CXCL11* (**Supplemental Figure 7**). Many of the genes reduced by BI-3812 and FX1 were also reduced by the presence of an NFκB inhibitor, suggesting that BCL6 acts on NFκB-responsive genes in endothelium; the functional effects of BCL6 BTB antagonism and NFκB inhibition might be similar. Additionally, a subset of genes that were not altered by TNFα treatment alone were upregulated by the combination of BTB domain inhibitors and TNFα together, such as *CD34∼, MMP1, RSAD2*, HMOX1*, LUCAT1*, AHR*, APLN*, CLDN5∼, EPAS1\** (*indicates BCL6 ChIP peak is present in gene, ∼ indicates ChIP peak near but not in gene).

The effect of BTB antagonists FX1 and BI-3812 on adhesion molecule expression was confirmed at the protein level by flow cytometry (**Figure 7b, 7c**). Both FX1 and BI-3812 significantly reduced TNFα-stimulated E-selectin and VCAM-1 cell surface expression at 4hr, and VCAM-1 at 24hr (**Figure 7b, 7c**). In contrast, the low potency FX1 analog 79-6 had no significant effect on any TNFα-induced adhesion molecules (**Figure 7d**). Conditioned supernatants from these experiments were tested for secreted chemokines and cytokines by multiplex assay (**Figure 7e**). FX1 strongly blocked production of G-CSF, GM-CSF, GROα, IL-6, IL-8, and MCP-1 by TNFα-treated endothelial cells at 4hr (**Figure 7e**). These results were highly concordant with effects on mRNA expression (**Supplemental Figure 7c, 7d**).

To link gene and protein expression with functional inflammation, we measured adherence of allogeneic PBMC to TNFα activated endothelial cells, in the absence or presence of BCL6 inhibitors, or an NFκB/NLRP3 inhibitor Bay11-7082 as a control. Concordant with suppression of adhesion molecule and chemokine expression, FX1 blocked PBMC adhesion to TNFα-activated endothelium at 4hr, with a similar inhibitory effect as NFκB inhibition with Bay11-7082 (**Figure 7e**). We used a second assay to confirm the effects, which showed that FX1 and BI-3812 substantially reduced the adhesion of U937 monocytes to TNFα-treated HAEC monolayers (**Figure 7f**).

These results suggest that NFκB and BCL6 may share regulated cistromes in endothelial cells, as previously reported in macrophages [14], which has important biological effects on endothelial adhesion molecule expression, chemokine production, and leukocyte adherence.

### Effect on NFκB Activation and Transcriptional Activity

Our next experiments focused on the functional actions of BCL6 in modifying NFκB-driven endothelial inflammation triggered by TNFα. First, we asked whether NFκB nuclear translocation was affected by BI-3802. As a control, pretreatment of cells with Bay11-7082 completely abrogated NFκB translocation into the nucleus after TNFα 30min. The NFκB nuclear signal was slightly augmented in the presence of BI-3802 after TNFα treatment for 30min and 60min (**Figure 8a**), BTB antagonists had no effect on NFκB nuclear localization at 30min and 60min (**Figure 8a**). We also tested TeloHAEC transduced with BCL6 shRNA or overexpression of BCL6, for TNFα-induced NFκB nuclear translocation. There was no difference in nuclear localization in response to TNFα (30min, 1hr, 3hr) under BCL6 knockdown.

**Figure 8.**
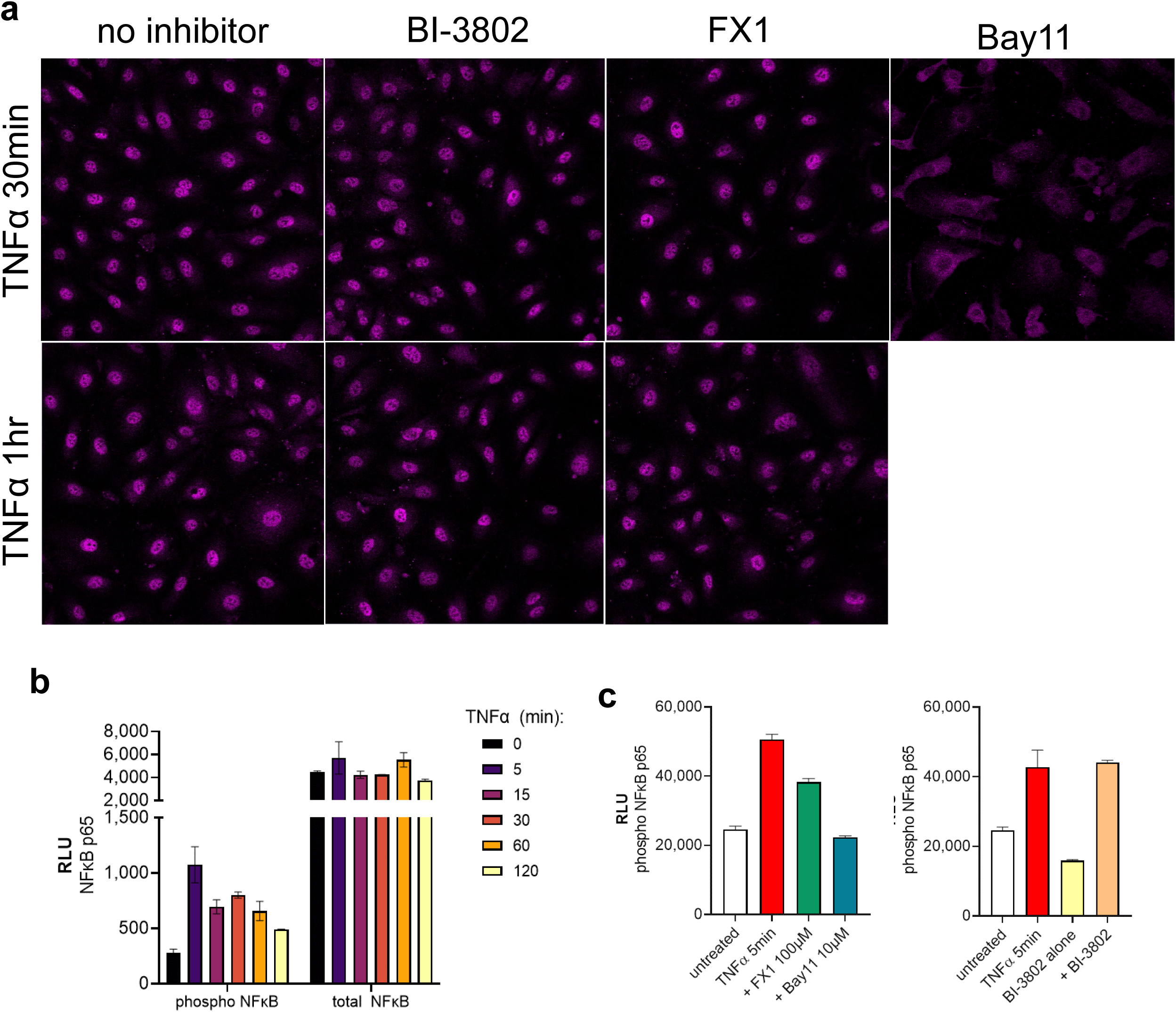
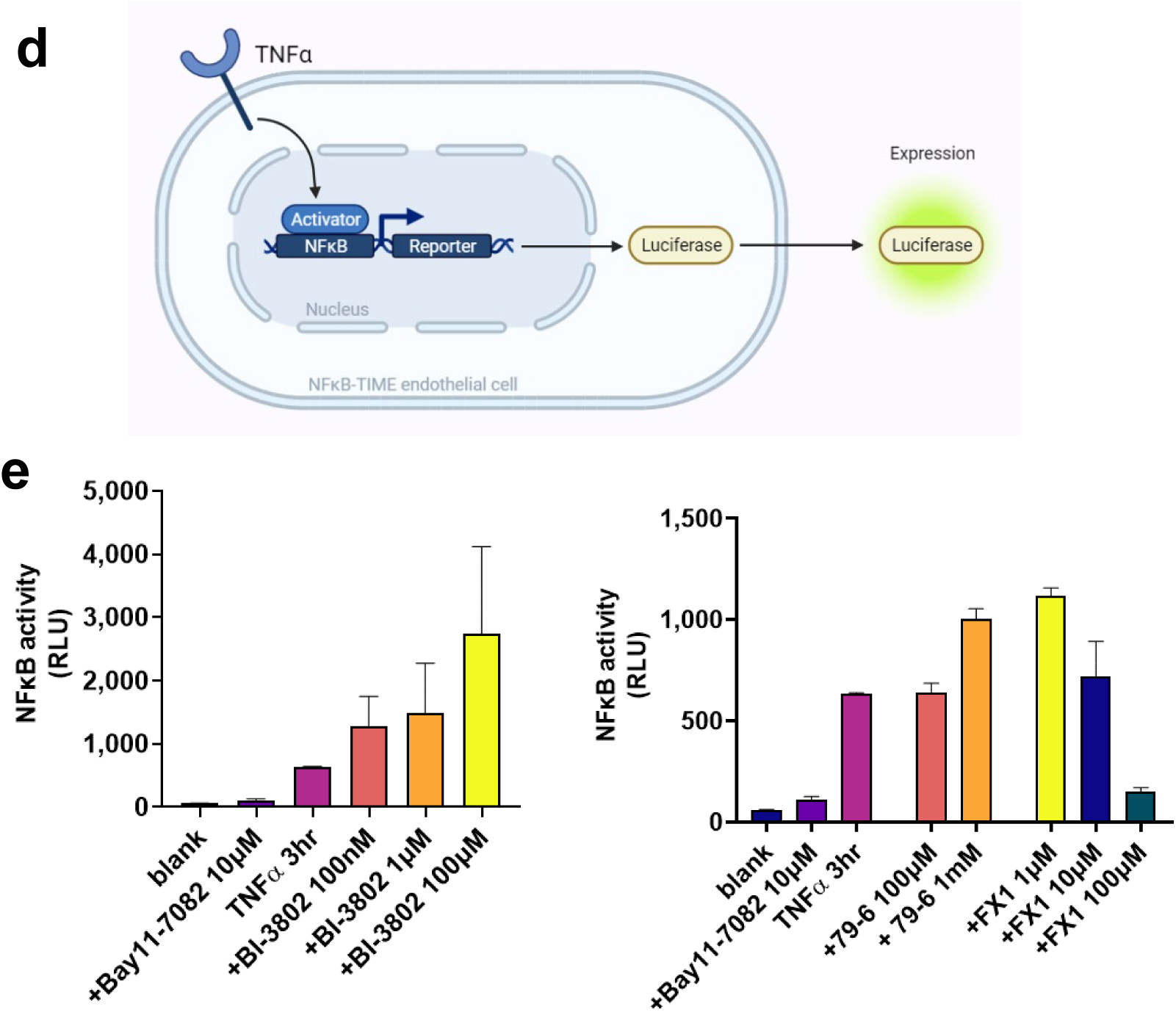
Effect of BTB inhibitors on NFκB signaling and activation. (a) NFKB intracellular localization in confluent HAEC. (b) Time course of NFKB phosphorylation (Ser536) measured by Lumit in HAEC (n=2 technical replicates). (c) Effect of FX1 (left panel) and BI-3802 (right panel) on NFKB phosphorylation. (d) NFKB-TIME immortalized endothelial cells express luciferase under the control of an NFKB responsive promoter. (e) NFKB-TIME cells were treated with TNFα in the presence of increasing doses of BI-3802 (left panel); or 79-6 or FX1 (right panel) and tested for NFKB transcriptional activity by luminescence. Results show average of two technical replicates, and are representative of 4 independent experiments.

Then, we measured activation of the NFκB pathway after 30min stimulation TNFα, in the presence of BCL6 pharmacological inhibitors. TNFα stimulation increased NFκB p65 phosphorylation at Ser536 within 5min, which was sustained to 2hr (**Figure 8b**). The NFκB inhibitor Bay11-7082 (10μM) prevented NFκB p65 phosphorylation, but neither the BTB domain inhibitor nor the degrading inhibitor BI-3802 affected phosphorylation of NFκB (p65 Ser536) (**Figure 8c**).

We next tested whether BCL6 directly affected TNFα-induced NFκB transcriptional activity in the luciferase reporter endothelial cell line NFKB-TIME (**Figure 8d**). Using this model, we previously showed that NFκB activity, as recorded by luciferase expression, increased by 2hr of TNFα exposure, peaking at 3hr, and declining but remaining above basal level at 24hr [75]. Bay11-7082 was used as a control and fully blocked NFκB activity (**Supplemental Figure 8**). BI-3802 increased the luciferase signal in a dose dependent manner, indicative of enhanced NFκB transcriptional activity. The low potency BTB antagonist 79-6 had no effect, but, contrary to our expectations and in line with the functional data above, FX1 strongly blocked NFκB-dependent transcription in a dose-dependent manner (1-100μM) (**Figure 8e**).

These results show that BCL6 affects NFκB transcriptional activity, but without affecting NFκB protein phosphorylation or nuclear localization under TNFα stimulation.

### Role for Histone Deacetylases

Interestingly, the effects of BTB domain inhibition and BCL6 depletion were not always opposite. For example, both BCL6 shRNA and BCL6-BTB antagonists reduced TNFα-induced *VCAM1*. We questioned whether the effects of BCL6 could be dependent on transcriptional repression via histone acetylation changes. We hypothesized that if the effects of BCL6 were HDAC dependent, then an HDAC inhibitor would mimic the suppression caused by BTB domain inhibitors FX1 and BI-3812. To address this hypothesis, HAEC were pre-treated with the HDACi TSA prior to stimulation with TNFα. Global gene expression changes were measured by RNA-Seq. Among genes that were highly induced by TNFα at 24hr, K means clustering showed two major groups of genes based on the response to TSA presence: approximately 70% of genes were augmented by TSA and the remaining predominantly suppressed by TSA (**Figure 9a**).

**Figure 9.**
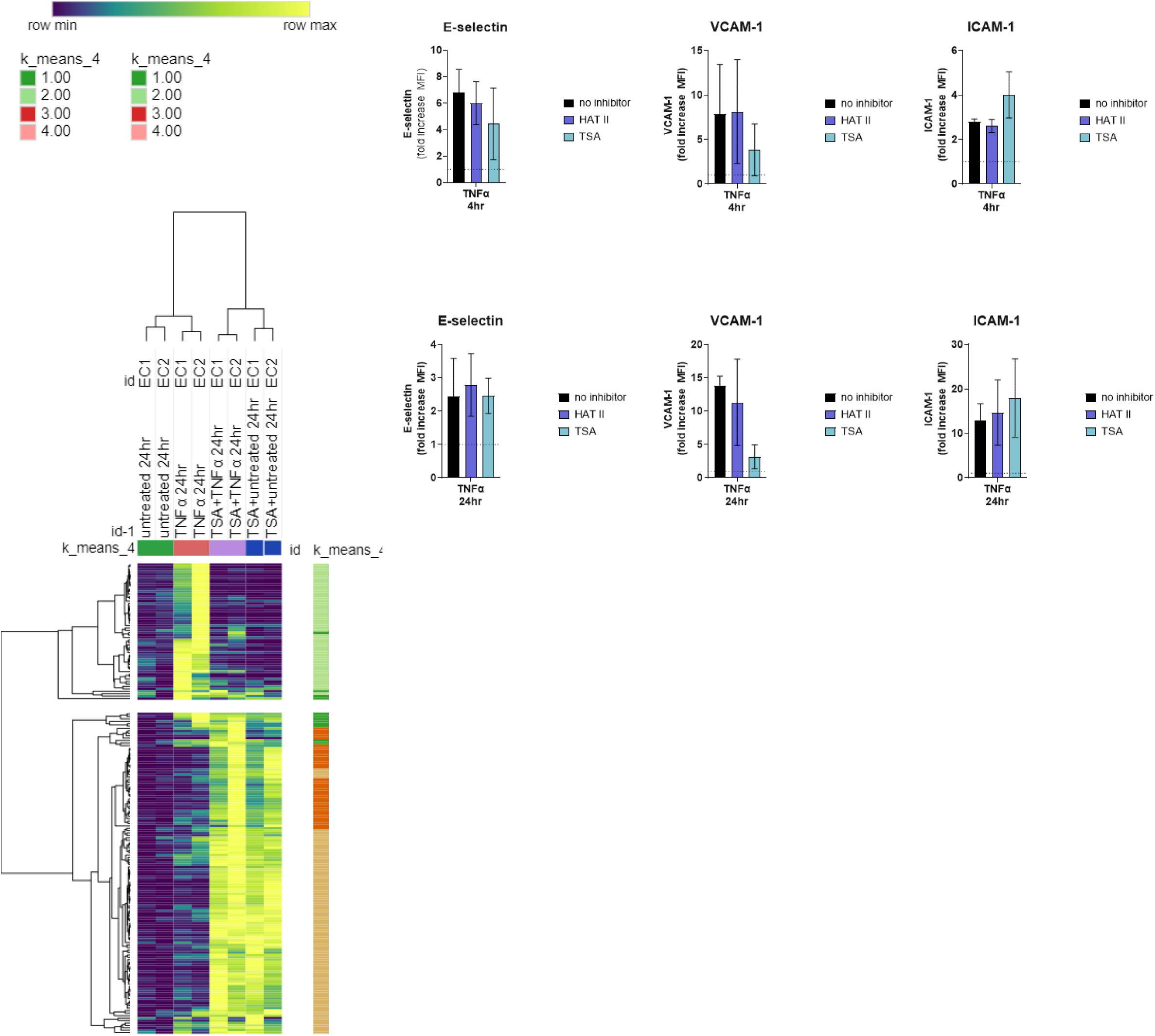
HDACs and endothelial pro-adhesive activation. Primary HAEC (n=2 biological replicates) were treated with the HDAC inhibitor TSA and stimulated with TNFα. (a) Gene expression was measured by RNA-Seq. Heat map shows relative expression of TNFα-induced genes. (b) Cell surface expression of adhesion molecules was measured by flow cytometry.

TNFα-induced expression of ICAM-1 and *CX3CL1* was not altered by HDACi, but *CCL5*, *CCL20*, and *BST2* were augmented by the presence of an HDACi at 24hr (**Supplemental Figure 9**). Although histone acetylation is a marker of active transcription, the majority of TNFα induced adhesion molecule and chemokine genes were suppressed by TSA, including *SELE*, *VCAM1*, and most other chemokines (*CXCL8*, *CCL2*, *CXCL1*, *CXCL2*, *CSF1*, *CXCL6*, *IL6*) (**Supplemental Figure 9**). We confirmed that TSA strongly suppressed TNFα-induced cell surface VCAM-1 expression at 4hr and 24hr, although E-selectin and ICAM-1 were not changed (**Figure 9b**). Differential effects were also seen for expression of NFκB-related genes. *NFKB1* at 4hr was partially inhibited by HDACi and totally inhibited at 24hr, while *BCL3*, *NFKB2* and *NFKBIB* were not affected, and *TNFAIP3*, *NFKBIA* and *NFKBIZ* were augmented by HDACi.

Genes suppressed by the HDAC inhibitor TSA showed a gain in H3K27Ac at or upstream of the transcriptional start site (TSS) under TNFα activation in human endothelium [66, 68, 69], including *VCAM1*, *SELE*, and *CXCL1*. However, other TNFα-induced genes that were unaffected or augmented by TSA did not exhibit changes in H3K27Ac, including *CCL5*, *CCL20*, *CX3CL1* and *BST2*.

### BCL6 repression of repressors

Since BCL6 antagonism affected more genes beyond its predicted direct targets, we examined the possibility of indirect regulatory actions. We examined microRNA expression in endothelial cells, stimulated with TNFα with and without BCL6 inhibitors. We found 6 microRNAs that were augmented by FX1: miR155, miR222, miR126, miR22, miR100 and miRlet7b (**Figure 10**). Further, we found highly significant BCL6 binding peaks in or near (within ±5kb) each of these miRNA genes (**Figure 10**), further supporting that BCL6 might directly regulate their expression. In contrast, no BCL6 binding peaks were found in other miRNAs that were expressed in endothelium but unaltered by BCL6 inhibition.

**Figure 10.**
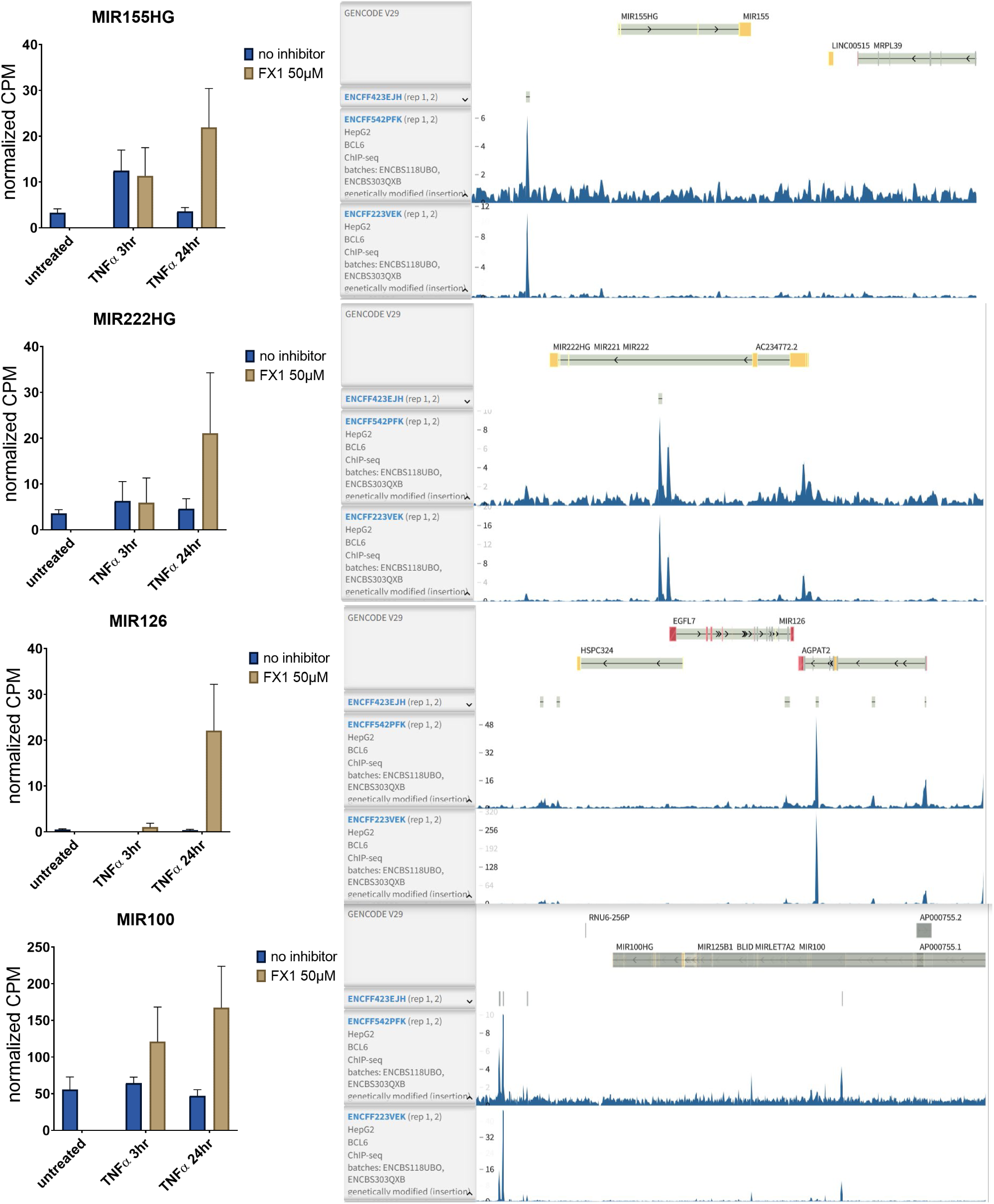
Effect of BTB inhibitors on miRNA expression in endothelial cells. Left panels show mRNA counts of MIR genes in HAEC (n=3) treated with FX1 and TNFα. Right panels show BCL6 ChIP-Seq tracks at each MIR gene from ENCODE.

Intriguingly, miR155 is already reported to be TNFα-inducible [76], and has complex functions in endothelial inflammation. In macrophages and B cells, BCL6 directly represses miR155 [77, 78]. Further, miR126 is known to target VCAM1 [79]. We lastly used miRgate [80] to identify targets of miR-126 and miR-155. Among endothelial inflammatory genes, miR-155 targets *CXCL5*, while miR-126 broadly targets chemokines and adhesion molecule genes, including *VCAM1, SELE, CXCL10, CXCL9, IL6* and *IL8* (but not *ICAM1* or *BST2*) (**Table 2**).

Based on these data, endothelial BCL6 can both directly regulate target genes and repress miRNAs. Therefore, inhibition of endothelial BCL6 would lead to an increase in miRNAs that prevent expression of NFκB target genes.

## Discussion

This study explored the role of BCL6 in endothelial cell inflammatory response and resolution of vascular inflammation. Focusing on TNFα stimulated endothelium, we found that NFκB RelA binding is increased to the BCL6 gene, and functionally BCL6 regulates endothelial adhesion molecule and chemokine expression. Taken together, our data support a previously undescribed role for BCL6 as a modifier of NFκB-dependent EC activation. Our working model is shown in **Figure 11**.

**Figure 11.**
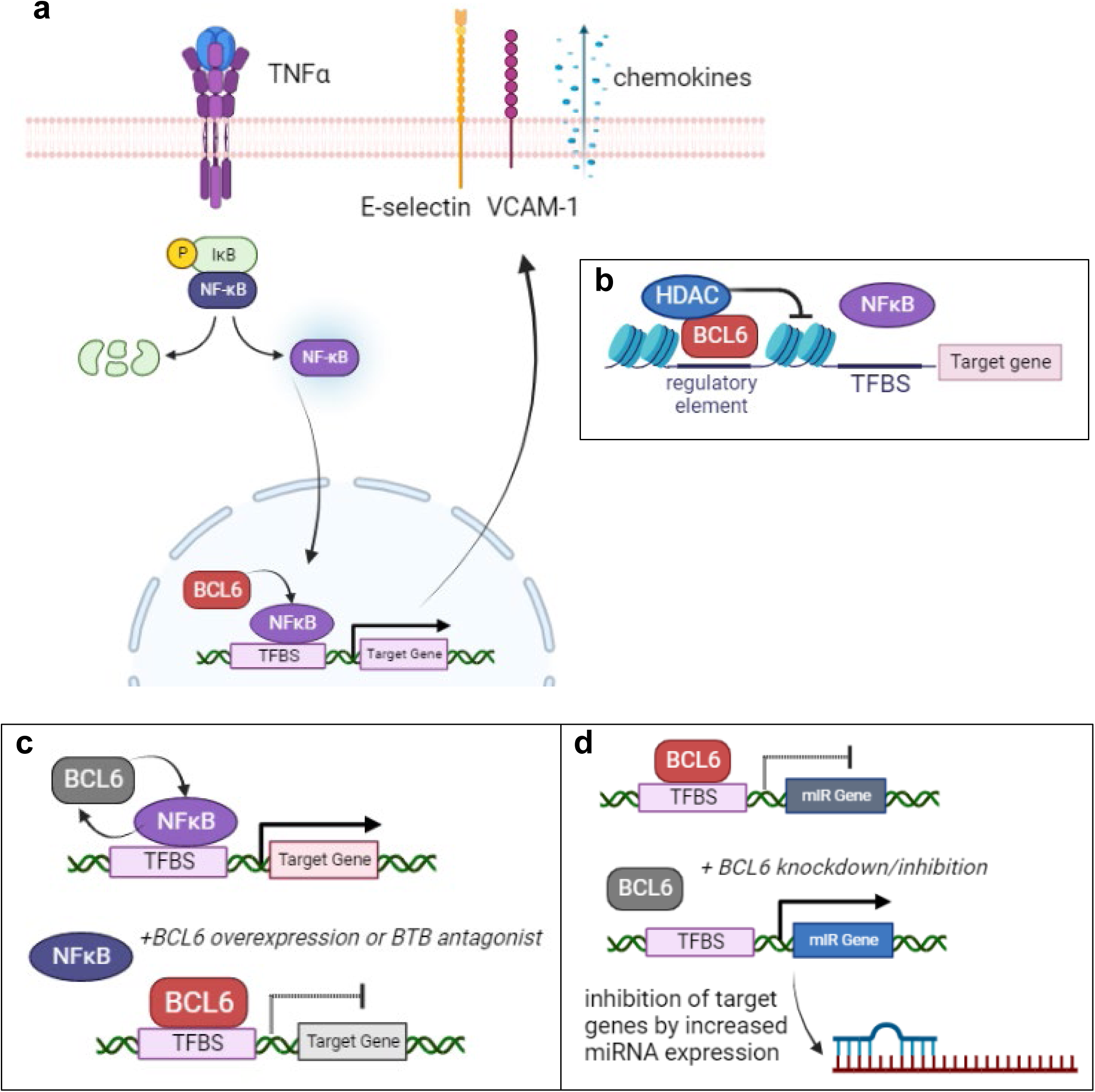
Proposed model of BCL6 and NFκB interactions in endothelial cells. (a) TNFα stimulation activates NFκB, which moves to the nucleus and displaces BCL6 at target genes, to activate expression of adhesion molecules and cytokines. (b) BCL6’s primary described mechanism of action is BTB-dependent association with HDACs, which suppress chromatin accessibility at target gnes. (c) An alternative mechanism of action is that BCL6 and NFκB cycle in their occupancy of inflammatory genes, and that BTB antagonists or overexpression of BCL6 enhances or sustains its occupancy at target genes, outcompeting NFκB to suppress gene expression. (d) A third potential mechanism of action is that BCL6 basally suppresses expression of miRNAs that regulate inflammatory genes. Loss of BCL6 by knockdown or BTB domain inhibition derepresses miRNAs, which in turn degrade their target genes, thereby reducing expression of inflammatory molecules. (figure created with biorender.com)

An interaction between BCL6 and NFκB has been reported in other cell types. For example, TNFα-induced NFκB activity and MCP-1 and IL-6 production were augmented in pancreatic cancer cells when BCL6 was knocked down, which increased monocyte invasion [81]. In B cell lymphoma, BCL6 repressed NFκB-activity in a manner that was partly dependent on DNA-binding (since a ZF mutant of BCL6 showed lower suppression of NFκB-luciferase than the full length) but potentially mediated by direct BCL6-NFκB binding [82]. In murine macrophages, BCL6 regulated approximately one quarter of the NFκB-controlled cistrome under LPS stimulation [14]. And, Gongol et al [83] reported that PARP1 suppresses BCL6 to promote vascular inflammation including enhanced VCAM1 expression and further demonstrated direct BCL6 binding to the VCAM1 promoter. Therefore, multiple potential mechanisms explain the effect of BCL6 on NFκB signaling.

Recently, an epigenetic comparison (H3K4me3 and H3K27me3) of blood vs. lymphatic dermal endothelium identified that significantly differentially DNA methylated regions were enriched for BCL6 binding motifs in blood endothelium [29]. Another prior study identified BCL6 DNA binding motifs among endothelial-restricted accessible chromatin [50].

Previous studies have demonstrated that cytokine stimulation of endothelial cells, including with TNFα, promotes histone modifications to establish latent enhancers [66, 68, 84] while repressing other genes. For example, H3K27Ac is increased at numerous genes in endothelial cells during TNFα activation [66]. Further, Inoue et al [85] showed that histone deacetylases were selectively required for TNFα-induced expression of VCAM-1 and fractalkine as well as maximal monocyte adhesion, although not for E-selectin or ICAM-1.

BCL6 has a reported role in remodeling of the chromatin landscape of target genes in non-vascular cells, in cooperation with corepressors that recruit HDAC3 [13, 34]. In B cells, knockdown of BCL6 enhanced H3K27Ac only at enhancers bound by both BCL6 and SMRT/NCoR, but not those occupied by BCL6 without SMRT [86]. Similarly, in the liver, loss of hepatocyte Bcl6 increased H3K27Ac at BCL6:SMRT:HDAC3 complex binding sites [35]. A mass spectrometric analysis of BCL6 binding proteins in SUDHL4 cells showed association with HDAC3, HDAC4, HDAC5, HDAC9 and Sin3A [32].

### Function in non-vascular cells

BCL6 was originally identified as a translocated oncogene in B cell lymphoma. In normal B cells, BCL6 activates B cell receptor and CD40 signal transduction by modulate MAPK and NF-kB pathways. Also, BCL6 repressed CD80 and CD274 affecting the B-T cell interactions. B cell differentiation is BCL6-influenced by modification in WNT-signaling and JAK/STAT. Its interactions with TGFβ affect late stages of germinal center (GC) differentiation modulating TGFβ receptors and transcription factors like PRDM1 and IRF4 [87].

Since these initial studies, it has become evident that BCL6 has broad functions in the immune system and beyond. In particular, BCL6 is a lineage-defining factor for T follicular helper cells [8], and systemic *Bcl6-/-* mice had aberrant Th2 and Th17 cells differentiation and pro-inflammatory cytokine expression. In fact, the absence of BCL6 repressed Th2 cell, favoring Th17 cell differentiation [88, 89]. Further, Global *Bcl6* deficient mice exhibit premature lethality due to dramatic myocarditis and pulmonary inflammation, which was not attributable to the leukocyte compartment [18, 90], which showed defective germinal center responses [17], but rather BCL6 in the heart was protective [18].

Thus, BCL6 is a critical transcription factor in T follicular helper cells involved in survival, differentiation, and migration. *In vivo* and *in vitro* assays demonstrated that BCL6 induced-TFH cell differentiation is not dependent of B cell, demonstrating that upregulation of BCL6 in CD4 T cells decrease PSGL1 expression. A deep genomic analysis established BCL6 as an effector of the Tfh linage. The report showed GC Tfh loci BCL6 binding sites at the promoter regions. These genes are associated with Tfh cell differentiation and migration, including *RORA, IFNGR1*, *STAT4*, *GATA3, PRDM1*, *S1PR4*, *CD69*, *LPP*, *FAIM3*, *PTEN*, *CASP8*, *FOXO3*. Interestingly, BCL6 binding to some elements of the STAT family (STAT1, STAT3, and STAT5) in T cells are related with wide genomic modifications directly involve in Tfh differentiation [8, 86, 90, 91].

In addition, BCL6 is expressed in macrophages, where it altered expression of genes-related inflammation, differentiation, cell adhesion, migration, and apoptosis. BCL6 represses the transcription of a subset of chemokine genes, like MCP-1, MCP-3, and MRP-1. MCP-1 promoter revealed three BCL-6 binding sites establishing cytokine interaction and regulation. Moreover, BCL6 binding flanks that of NFκB p65 in a reciprocal regulation of the inflammatory response. KLF6 a pro-inflammatory transcription factor, suppressed Bcl6 expression and promoted pro-inflammatory phenotype by PRDM1. Blockade of BCL6 in KLF6-deficient macrophages reversed attenuation of pro-inflammatory chemokine/cytokine expression, cellular migration, and invasion [14, 92–94]. In macrophages participating in atherosclerosis, BCL6 is targeted by AngII, which decreases its ability to repress inflammatory genes. This mechanism is mediated by angiotensin type 1 (AT-1) receptor.

miR-155 mediates macrophages inflammatory response in atherosclerosis. The miR-155 actions on macrophage efferocytosis are mediated by Bcl6. In fact, the suppression of *Bcl6* by miR-155 increased RhoA activity and lowered efferocytosis. Another study demonstrated that miR-155 promoted macrophages activation by antagonizing BCL6-mediated inhibition of NFκB and reduced atherogenic cytokine expression.[25, 78, 93]. In neutrophils, the ablation of BCL6 gene resulted in decreased inflammatory cytokines and promotion of TNFα signaling via NFκB gene. In broncho alveolar lavage fluid, BCL6-knockout-neutrophil upregulated apoptotic genes including Casp 3, PmiP1 and Pdcd4 suggesting a BCL6 active participation in neutrophil survival post-influenza A virus infection in the lungs [95].

Outside the immune system, BCL6 functions in adipocytes as a regulator of glucose metabolism and lipid synthesis. BCL6 abrogation in these cells improved insulin sensitivity and lipogenesis. Knockout mice exhibited more adiposity, and tissue inflammation [74]; and in their livers, the analysis of the BCL6 cistrome demonstrated response elements for lipid and ketone metabolism, and PPAR signaling. DNA motif prediction analysis resulted in PPARs colocalizations in BCL6 genes specially under fasting challenge. In the liver, BCL6 broadly regulates metabolic and oncogenic processes [[35, 96, 97]], and suppresses growth hormone signaling, insulin resistance and steatosis in an adipocyte-intrinsic manner [74, 98]. Furthermore, BCL6 depletion reduced the effect of metabolic impairment associated to PPARα deficiency reflected in a dysregulated fatty acid oxidation, hypo-ketonemia and steatosis sensitivity. Other BCL6 liver knockout studies reported lower hepatic levels of triglyceride contrary to serum cholesterol, and HDL cholesterol. Moreover, BCL6 are associated with metabolic regulation thought upregulated β-oxidation-related genes and Socs2 as well as downregulated proinflammatory cytokines like TNFα and IL-6 [[35, 96, 97].

### Function in vascular cells

Other evidence that BCL6 is an effector of vascular smooth muscle established an opposite regulation with AngII. BCL6 overexpression inhibited VSMC proliferation by reducing NOX4 expression, and oxidative stress by lower NADPH oxidase activity and ROS production [99]. Reports of BCL6 antagonizing NOTCH activation have established its negative effects on vessels sprouting formation and proliferation [30].

Intriguingly, BCL6 was among the transcription factor motifs whose variants affected ERG binding in HAEC, suggesting that BCL6 might overlap with endothelial master regulators [65]. Other reports demonstrate BCL6 antagonizing NOTCH activation [30]. Further, BCL6 suppresses VCAM1, MCP1 and MCP3 under AMPK activation in endothelium with demonstrated direct binding to the promoter of these genes, presumably through an adaptive stress response since BCL6 binding was increased at the promoters under pulsatile shear flow [83]. In endothelium, BCL6 is possibly sequestered by PPARδ, which when activated releases BCL6, BCL6 increases at the promoter of VCAM1 and represses its expression by TNFα; therefore BCL6 carries out the repressive function of PPAR agonists [31]. This direct interaction between PPAR and BCL6 is similar to that in macrophages, where repressive/inactive PPAR binds to BCL6, at a similar binding domain as HDACs, and upon dissociation of PPAR-BCL6, BCL6 can repress gene expression [100]. An interaction between BCL6 and PPAR was also described in the liver, although here through proximal binding at genomic regulatory sites [35].

### Potential for BCL6 to ameliorate vascular inflammation

Surprisingly, we found that inhibitors that block complex formation with BCOR/NCOR and presumably also prevent HDAC3 recruitment, instead suppress a majority of TNF-induced gene expression. This was consistent with a paper that showed that one of these inhibitors, FX1, enhanced BCL6 binding to target genes [41]. Further, treatment with the BTB inhibitor FX1 suppressed tissue inflammation in several mouse models, contrary to expectations that BTB-dependent functions were anti-inflammatory [47, 48]. These reports are in line with our *in vitro* data showing a potent dampening effect of FX1 and BI-3812 on TNFα-induced gene expression. We had expected that antagonism of BCL6 association with corepressors would relieve gene repression and exacerbate inflammatory responses. Thus, an alternative hypothesis is that BCL6 has a BTB-independent function to antagonize NFκB, for example by competitive blocking of NFκB or through repression of EP300 as in DBLCL [101]. Another explanation could be that BCL6 is induced because it is required to antagonize anti-inflammatory genes and permit TNF induced gene expression, and so impaired BCL6 function enhances the negative regulatory effects.

BCL6 KO T-cells or pharmacological BCL6 inhibitors in a chronic graft-versus-host disease (cGVHD) murine model demonstrated improve pulmonary function by reduce GC B-cell and TFH cell frequencies, and pulmonary collagen and immunoglobulins deposition [102]. In a renal hypertension model, BCL6 reduced NLRP3 and IL-1β expression [103]; however, we did not observe NLRP3 expression under basal or TNFα stimulated conditions in endothelium. In a murine sepsis model, the BCL6 BTB inhibitor FX1, improved survival, attenuated serum inflammatory cytokines and monocytes/macrophages infiltration. FX1 downregulated the expression of TNF-α, IL-1β, IL-6 and MCP-1, and the activation of NF-KB by enforce BCL6 binding to target promoters [48]. Wound healing was improved by the absence of miR-155. Stimuli with IL-4 promote miR-155 targets like BCL6 suggesting its participation in healing process [104].

Although suppression of many TNFα-induced genes by BTB inhibitors was initially unexpected, our observations are in line with prior findings that 79-6 and FX1 attenuate inflammation in a murine sepsis model and chronic graft versus host disease [102], potentially due to augmented BCL6 binding to DNA [41, 48] and inferred enhancement of its repressive activity.

Based on the assumption that BCL6 is a transcription repressor, it was somewhat unexpected that treatment of mice with BTB inhibitors diminished rather than enhanced inflammation and organ damage in mouse models [48, 102]. A proposed explanation is that BTB inhibitors, while blocking association with BCOR and NCOR/SMRT, in fact augment BCL6 binding to DNA, as has been shown in other cells [41], suggesting a co-repressor independent function of transcriptional regulation by BCL6. This mechanistic model would explain why BTB antagonists repress select NFκB target genes in our results, by increasing BCL6 occupancy and outcompeting or otherwise inhibiting NFκB activity (**Figure 11b**).

Surprisingly, impairment of the BTB domain through which corepressors bind did not universally impair BCL6 repressive activity, at least in macrophages[105]. Moreover, mice expressing a BTB mutant of *Bcl6* did not exhibit the profound inflammatory phenotype seen in systemic *Bcl6* knockouts [105]. Importantly, mutation or antagonism of the BTB domain does not reduce binding of BCL6 to its DNA targets and indeed may enhance binding [41, 48]. In hepatocytes and macrophages, roughly 20% of the BCL6 genomic binding sites were unique to BCL6 and did not overlap with SMRT or NCOR[14, 35], and in B cells, H3K27Ac at BCL6-bound enhancers lacking SMRT, was not affected by 79-6, a BTB domain inhibitor [36]. Our data indicates that BCL6 perturbation suppresses more than half of TNFα-responsive genes in endothelium. Given the intersection of BCL6 and NFκB DNA binding sequences, we hypothesize that BCL6 directly represses NFκB target gene expression in a corepressor-independent manner, as shown for AMPK-induced VCAM-1 and MCP-1 [31]. Surprisingly, BCL6 inhibitors affected genes that did not have demonstrable BCL6 binding, at least in untreated HepG2 cells. It is possible that BCL6 affects these genes through secondary or tertiary effects; and/or that BCL6 binding patterns are altered in endothelium after TNFα treatment. Experiments are planned to define the BCL6 cistrome in endothelium at baseline and under TNFα activation.

Possible explanations for our findings that BTB domain inhibitors suppressed a significant proportion of TNFα-induced inflammatory genes are that 1) BCL6 exhibits independent anti-inflammatory functions, enhanced by these inhibitors; or 2) conversely that that BCL6 corepressor complex is required to suppress basally expressed quiescence genes to promote pro-inflammatory function. Our data show that BCL6 BTB inhibitors suppress TNFα-induced miR-155 (**Fig 10**), a TNFα inducible microRNA that regulates endothelial adhesion molecule expression [106, 107]. There are established reciprocal interactions between BCL6 and miR-155HG [77, 78, 108]. BCL6 may actively repress genes like miR155 that antagonize TNFα activation, to secondarily dampen inflammation (**Figure 11d**). Therefore, BCL6 likely performs transcriptional modification that is selective and context dependent. Studies are ongoing to decipher how BCL6 reshapes the NFκB-driven endothelial response, by parsing two putative mechanisms: *i)* corepressor-dependent epigenetic silencing and *ii)* corepressor-independent modification of NFκB target genes.

### Chromatin Remodeling in endothelial inflammation

TNFα elicits a multi-phase response that begins with immediate early genes with more accessible chromatin, and transitions to late phase genes after chromatin remodeling [109–112]. TNFα treatment also dramatically alters the chromatin state in endothelium, with activation of latent enhancers at inflammatory response genes and repression of basal superenhancers [68, 113]. Inhibition of HDAC3 by Trichostatin A (TSA) in EC abrogated TNFα-induced *VCAM1* and *CX3CL1*, but not *CXCL1*/GROα, ICAM-1 or E-selectin expression [85], highlighting distinct requirements for H3K27Ac chromatin restructuring of endothelial NFκB targets. These data provide a rationale for investigating the epigenetic effect of BCL6 in NFκB-mediated transcription.

Histone acetyltransferase p300 is rapidly recruited to the promoter of NFκB target genes under TNFα stimulation, such as *SELE*, with concordant nucleosome remodeling including an increase in 5’ diacetylated H3 and H3K9 acetylation at the promoter and CDS [Edelstein JBC 2005 PMID 15671023]. In particular, HDAC2 was specifically bound to the cytokine response region upstream of the TSS, along with the scaffold protein Sin3a, by 30min while HDAC3 assembles downstream of the TSS slightly later at 60min.

HDAC substrates include more than just histones. For example, HDAC9 deacetylates IKKα and IKKβ to inhibit their activity and thereby increase activation/phosphorylation of NFκB [114]. Indeed, knockdown of HDAC9 or inhibition with TMP195 reduced NFκB-dependent vascular inflammatory responses. Similarly, HDAC3 reportedly removes inhibitory deacetylases from p65 NFκB [115]. In our hands, TSA had no effect on TNFα-induced NFκB phosphorylation (data not shown), but did substantially block HDAC activity in the reporter assay as expected.

### Limitations

An important limitation of our study was that experiments were predominantly conducted in *in vitro* cultures of endothelial cells. Follow-up work is ongoing to further examine the *in vivo* effects of BCL6 in regulating vascular inflammation. We are also actively investigating the overlap between the BCL6 and NFκB cistromes, to define the mechanisms by which BCL6 controls NFκB-dependent transcription.

### Summary

Chronic vascular inflammation underlies the pathogenesis of many diseases, yet there is a dearth of therapies to dampen endothelial response to inflammation and injury. Despite the surge of interest in endothelial cell heterogeneity and its ramifications on human health, fundamental questions remain about how differential gene expression translates to organotypic vascular homeostasis and inflammation. These data expand our understanding of basic endothelial cell biology, providing insight into the function of the conserved transcriptional regulator BCL6 in endothelial cell homeostasis and inflammation.

## Acknowledgements

The authors would like to extend their thanks and acknowledge the technical contributions from the following individuals: the UCLA Immune Assessment Core, including E. Cho, M. Rossetti, G. Sunga, and M. Cappelletti, for technical assistance in performing the Luminex assays and development of flow cytometry panels; the UCLA Center for Systems Biomedicine, particularly to E. Faure for performance of Nanostring assays; to the UCLA Translational Pathology Core Laboratory, particularly to Y. Li for immunofluorescence staining of human tissue; the UCLA Technology Center for Genomics and Bioinformatics, especially to X. Li for RNA-Sequencing; and to L. Cerchietti, A. Melnick and M. Koegl for conceptual discussions on BCL6 inhibitors and mechanism of action.

## Funding

This work was supported in part by the Norman E. Shumway Career Development Award from the International Society for Heart and Lung Transplantation and Enduring Hearts (to NMV); the UCLA Faculty Development Award (to NMV); the Faculty Development Research Grant from the American Society of Transplantation (to NMV); and the National Institutes of Health R01-AI135201 01A1 (ER), U01-AI136816 03 (JM), and U01-AI163086 01 (JM/RH).

## Disclosures

None.

---

**Supplemental Figure 1.**
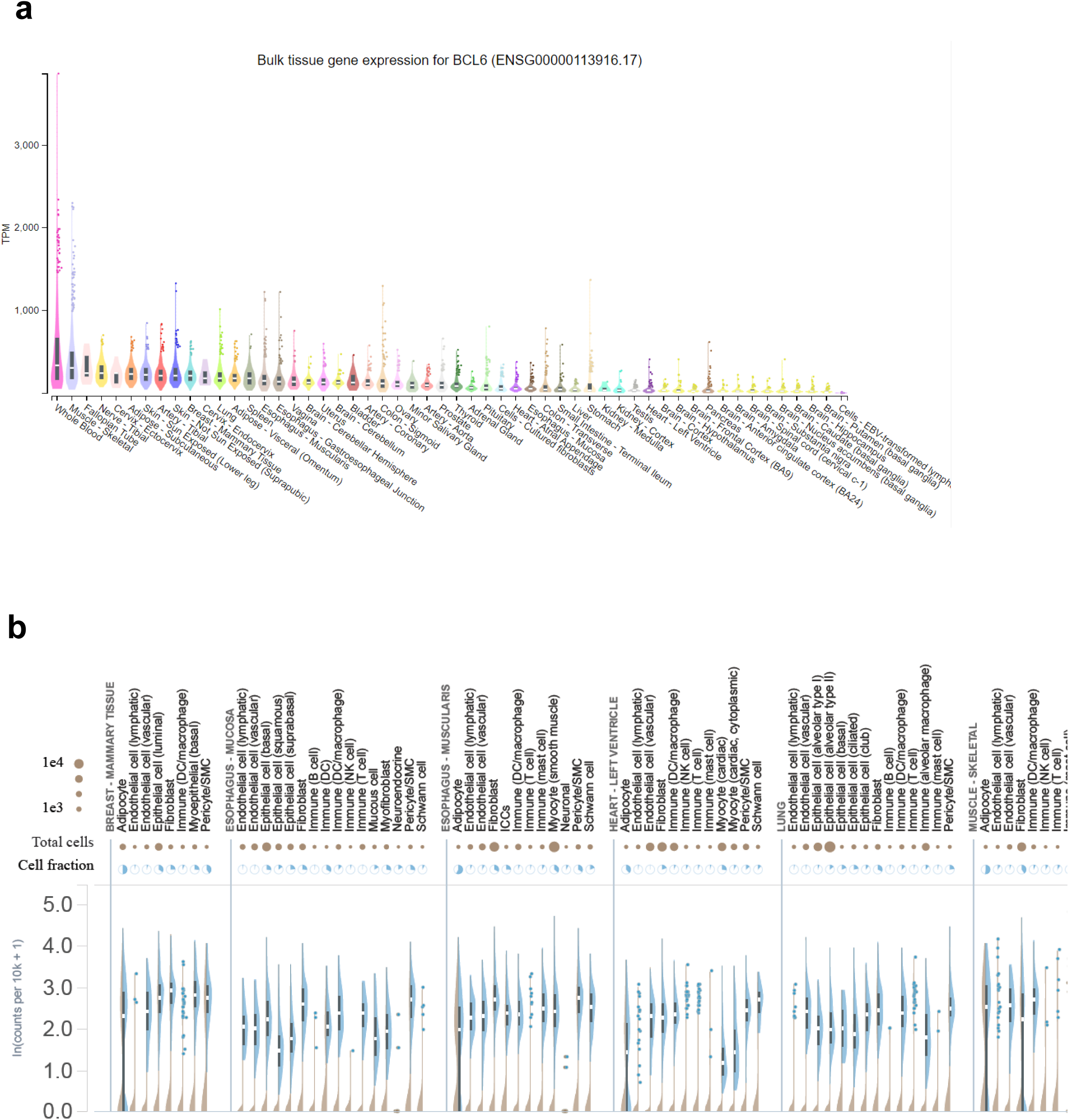
RNA expression levels of BCL6 in human endothelial cells from GTEX. (a) bulk gene expression (b) snRNA-Seq

**Supplemental Figure 2.**
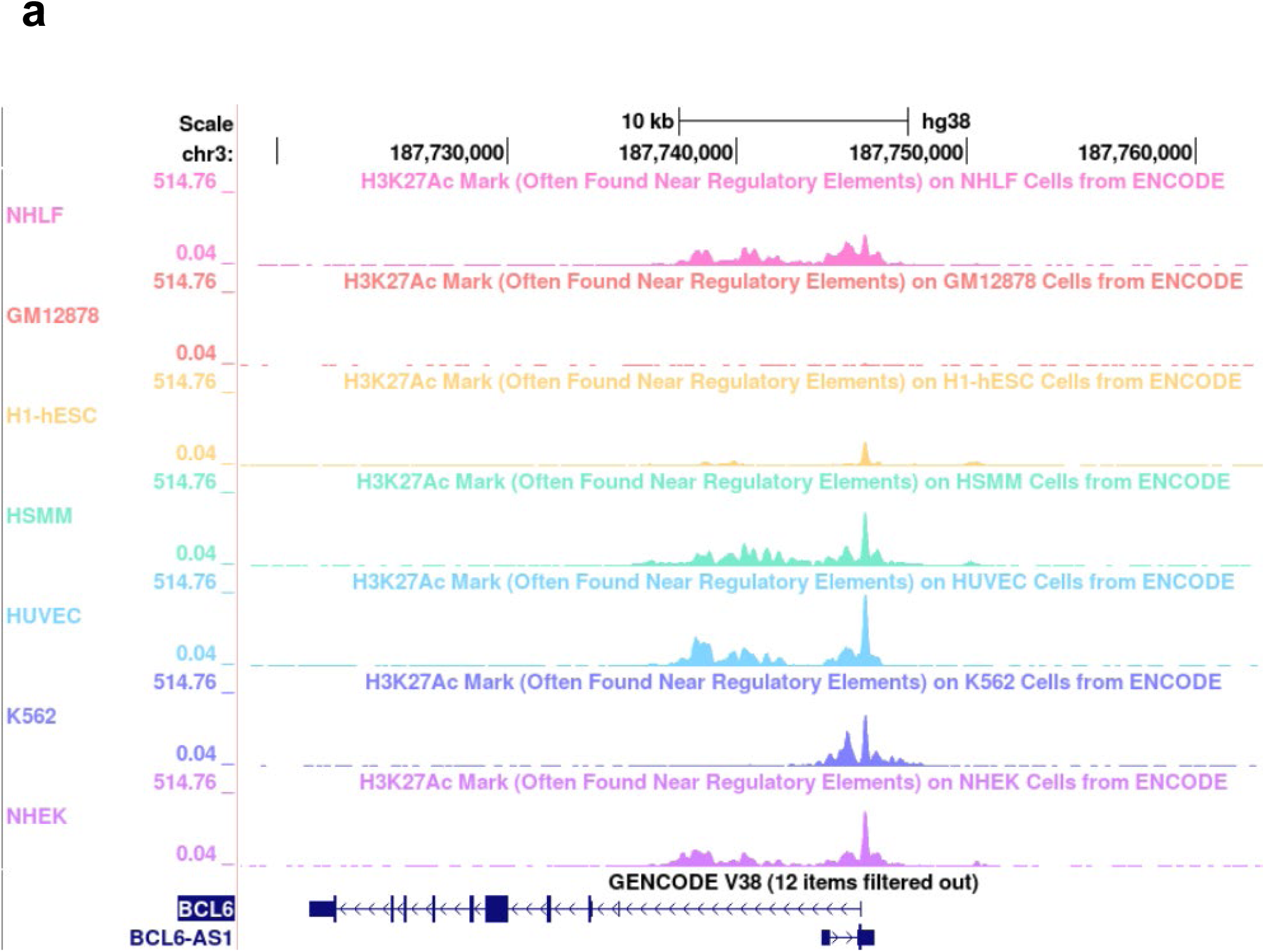
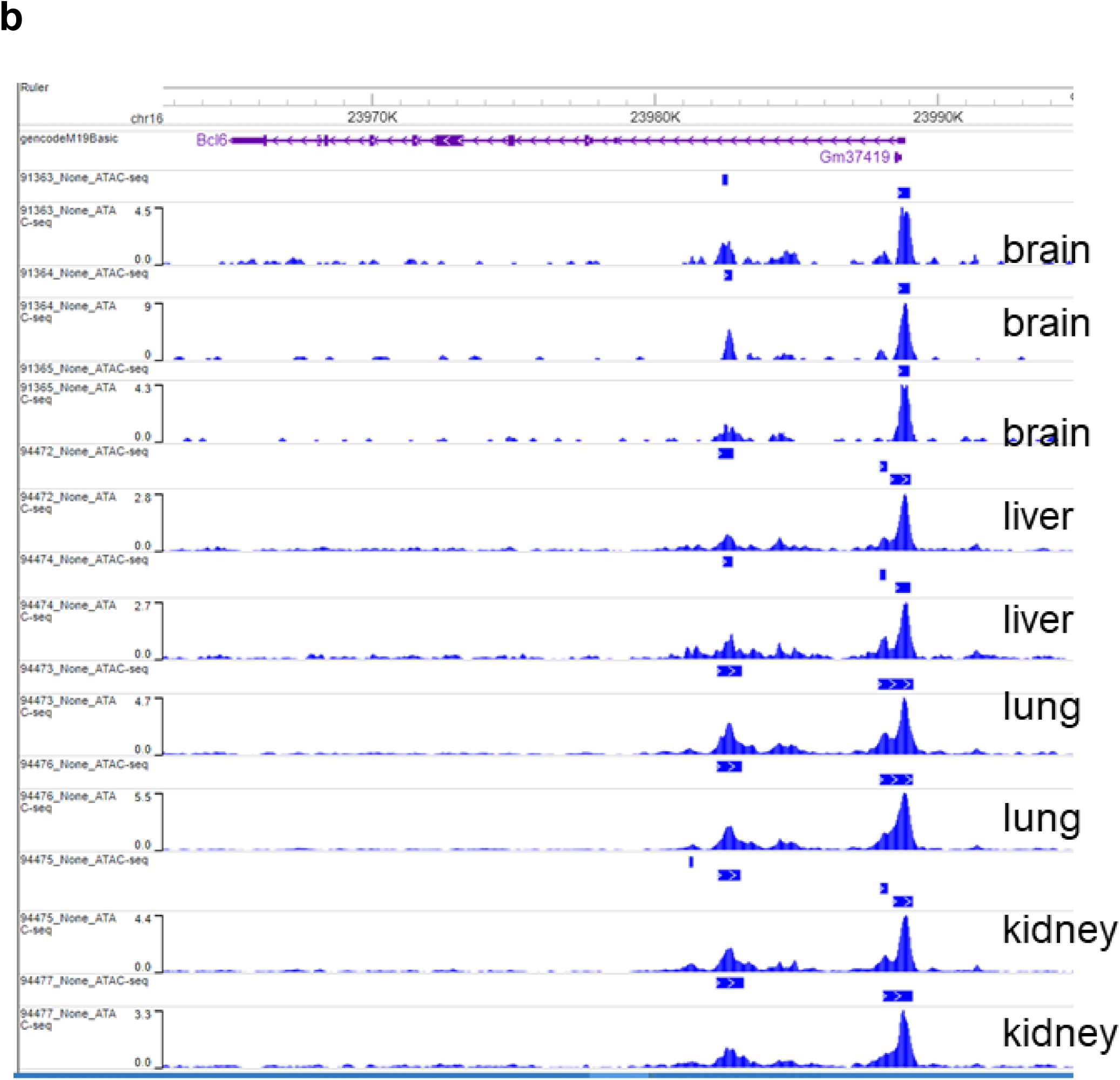
(a) BCL6 gene shows high chromatin accessibility in many cell types, including in endothelial cells. ENCODE tracks for H3K27Ac ChIP-Seq in 7 different human cell types. BCL6 gene shows high chromatin accessibility in many cell endothelial cells from different murine organs. (b) ATAC-Seq tracks from the Vascular Endothelial Cell Trans-omics Resource Database (Sabbagh) in endothelial cells isolated from the mouse brain, liver, lung and kidney.

**Supplemental Figure 3.**
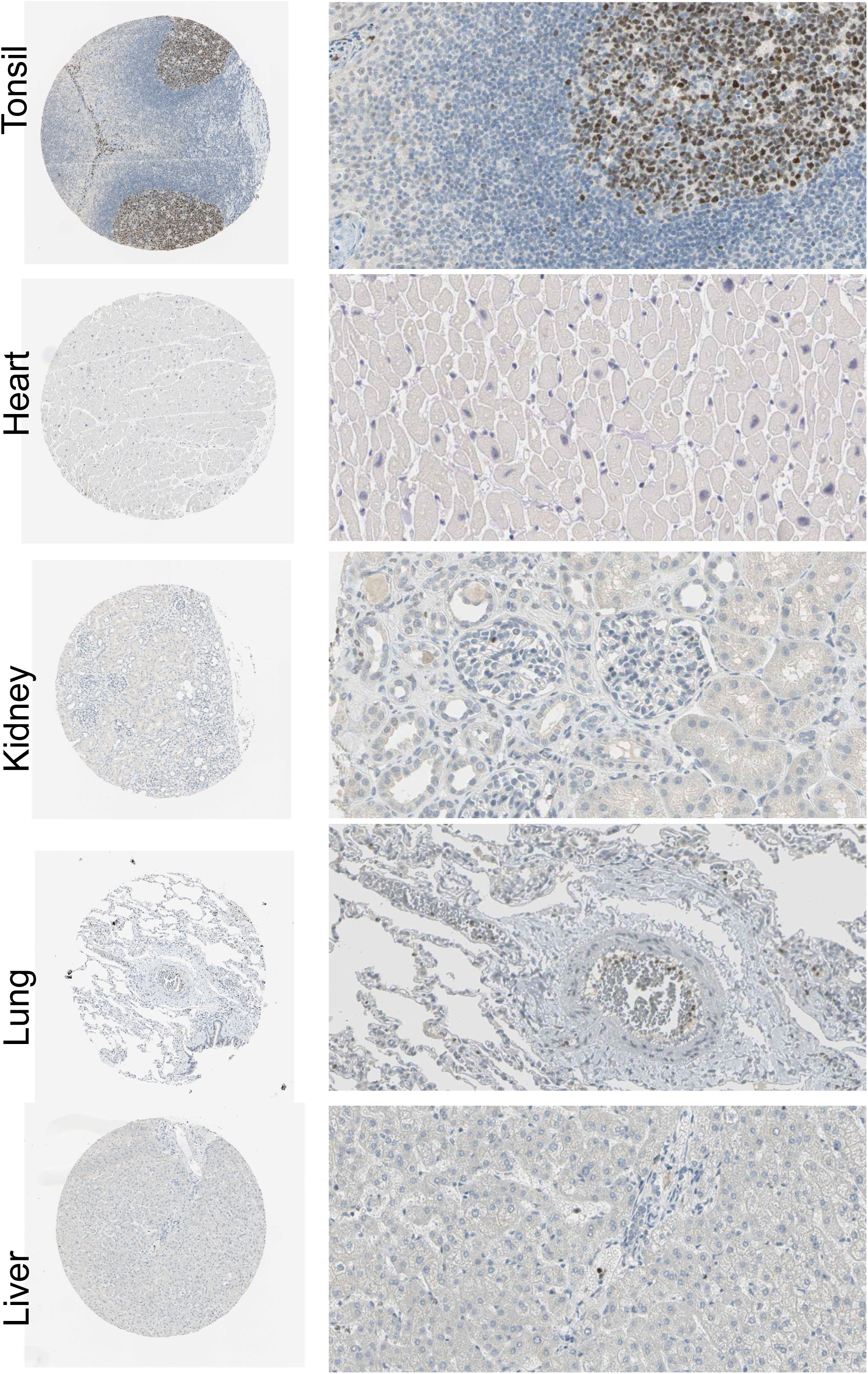
BCL6 protein expression in normal human heart, lung, liver and kidney from Human Protein Atlas.

**Supplemental Figure 4.**
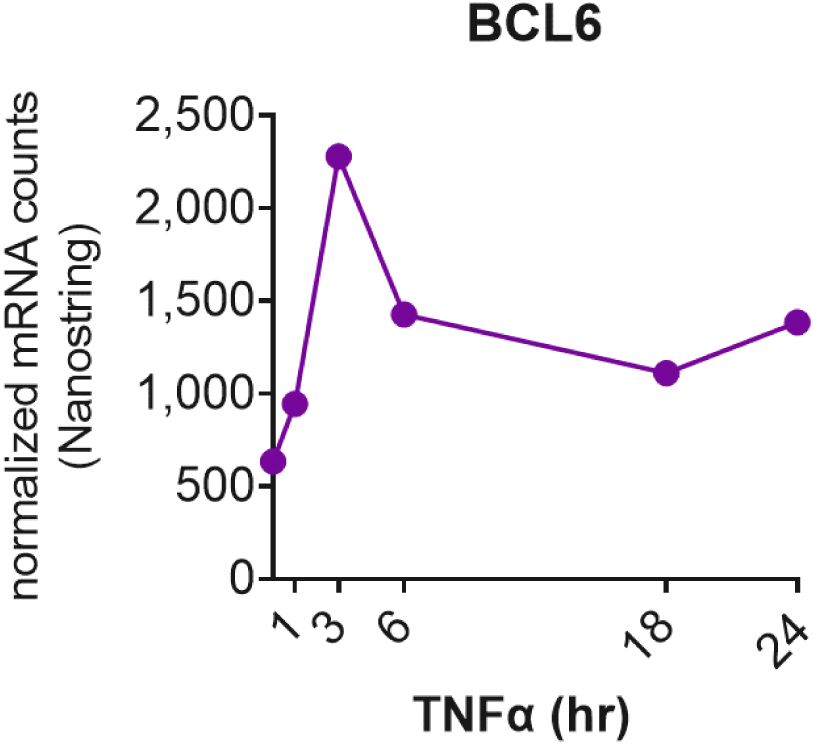
(a) Time course of BCL6 expression. HMEC-1 endothelial cells were treated for 1, 3, 6, 18 and 24hr, and mRNA expression as tested by Nanostring.

**Supplemental Figure 5.**
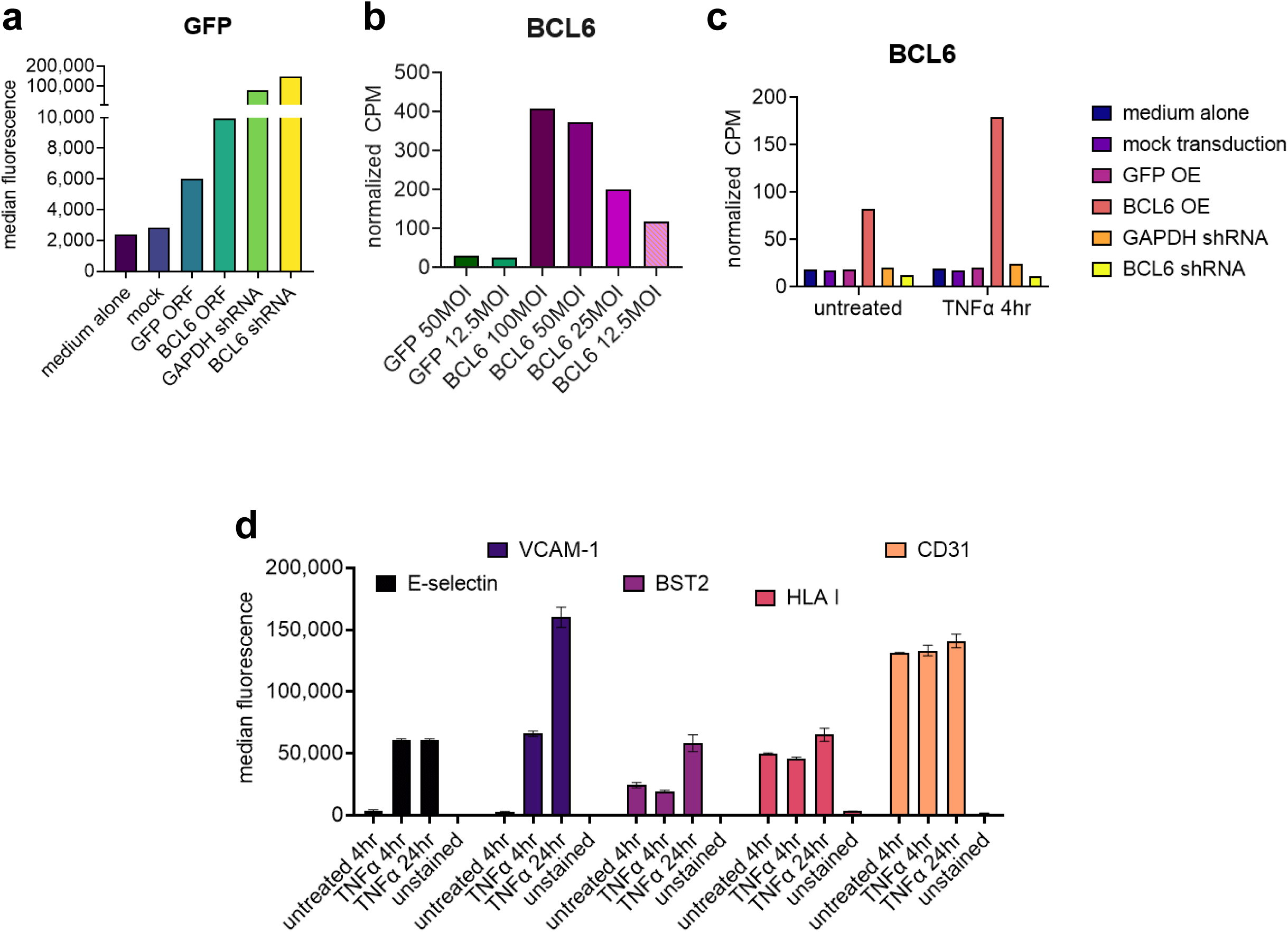
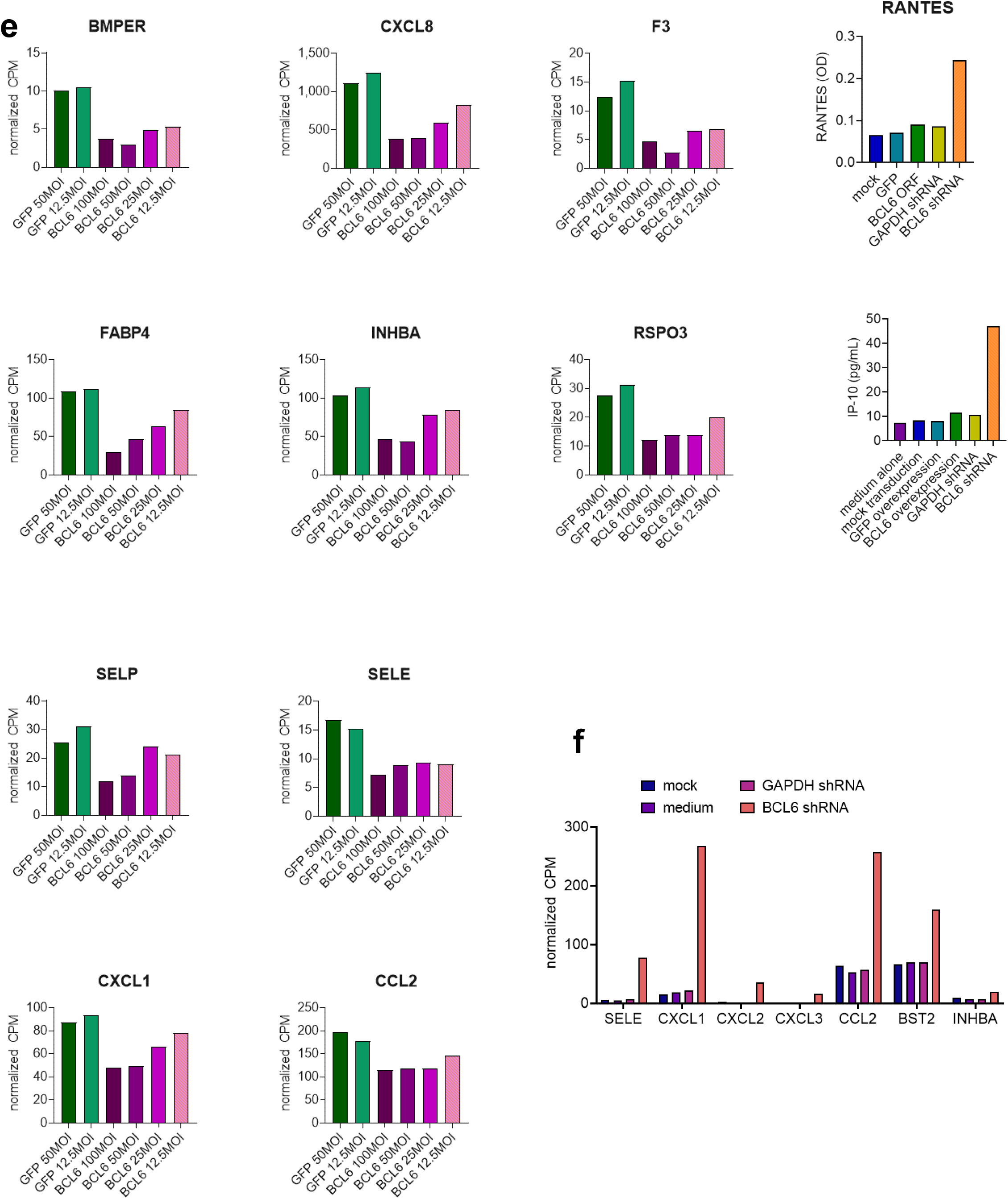
Optimization of lentivirus knockdown and overexpression of BCL6 in endothelial cells. (a) GFP reporter expression measured by flow cytometr in lentivirus-transduced TeloHAEC. (b) mRNA normalized CPM of BCL6 transcript in primary HAEC transduced with increasing MOI of lentivirus expressing either GFP alone or the BCL6 open reading frame (left panel), or TeloHAEC transduced with 50 MOI of shRNA or overexpression vector (right panel). (c) TeloHAEC respond to TNFα by upregulating canonical adhesion molecules. TeloHAEC were treated with TNFα for 4hr and 24hr, and expression of cell surface adhesion molecules and LA class I was measured by multiparameter flow cytometry (n=2 technical replicates). Impact of BCL6 overexpression and knockdown on basal gene expression in endothelial cells.

**Supplemental Figure 6.**
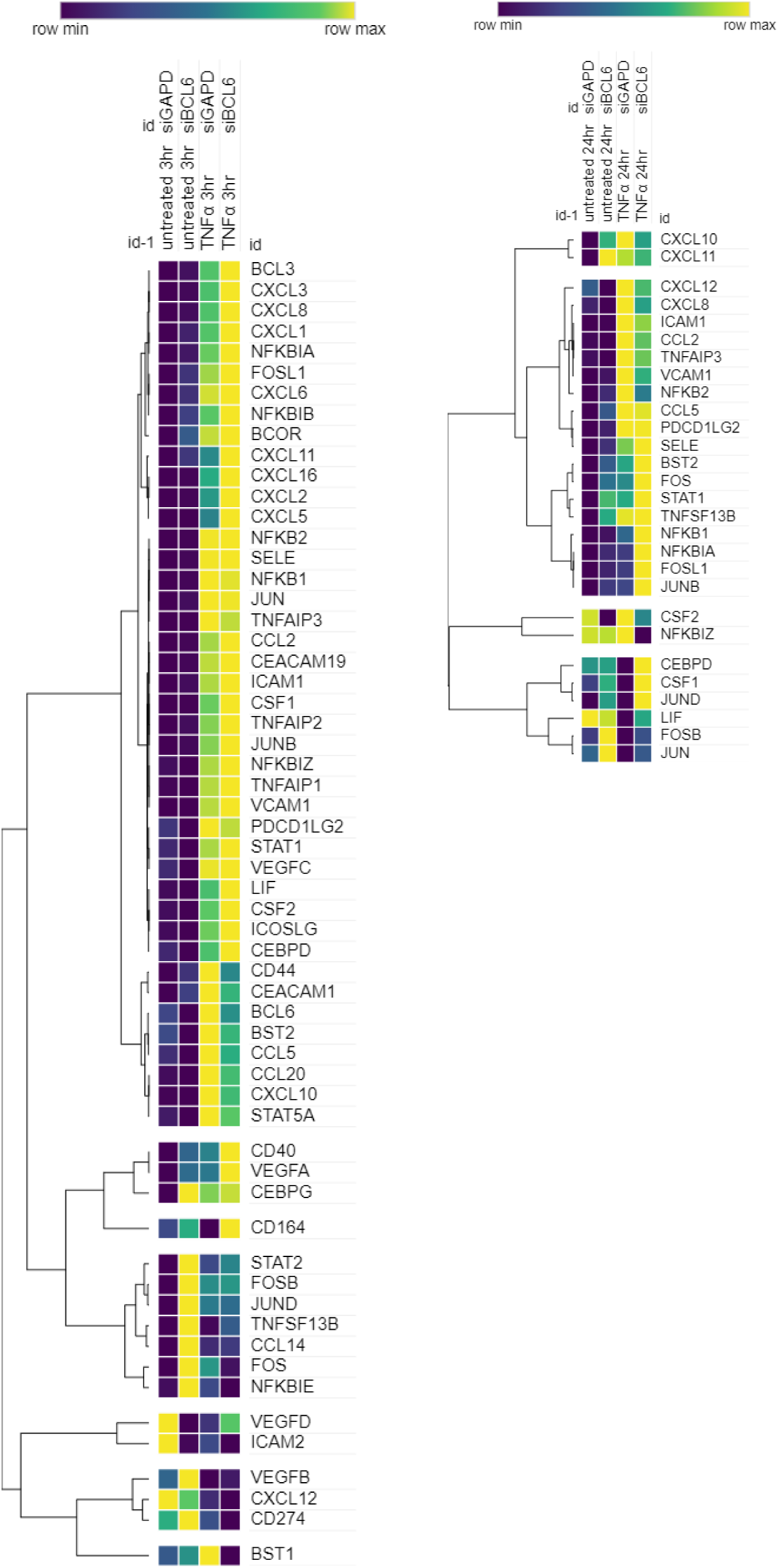
siRNA knockdown of BCL6 in primary cells only achieved about a 50% decrease in BCL6. Nevertheless, siRNA to BCL6 did affect expression of TNFα-induced genes, in line with shRNA in immortalized endothelium.

**Supplemental Figure 7.**
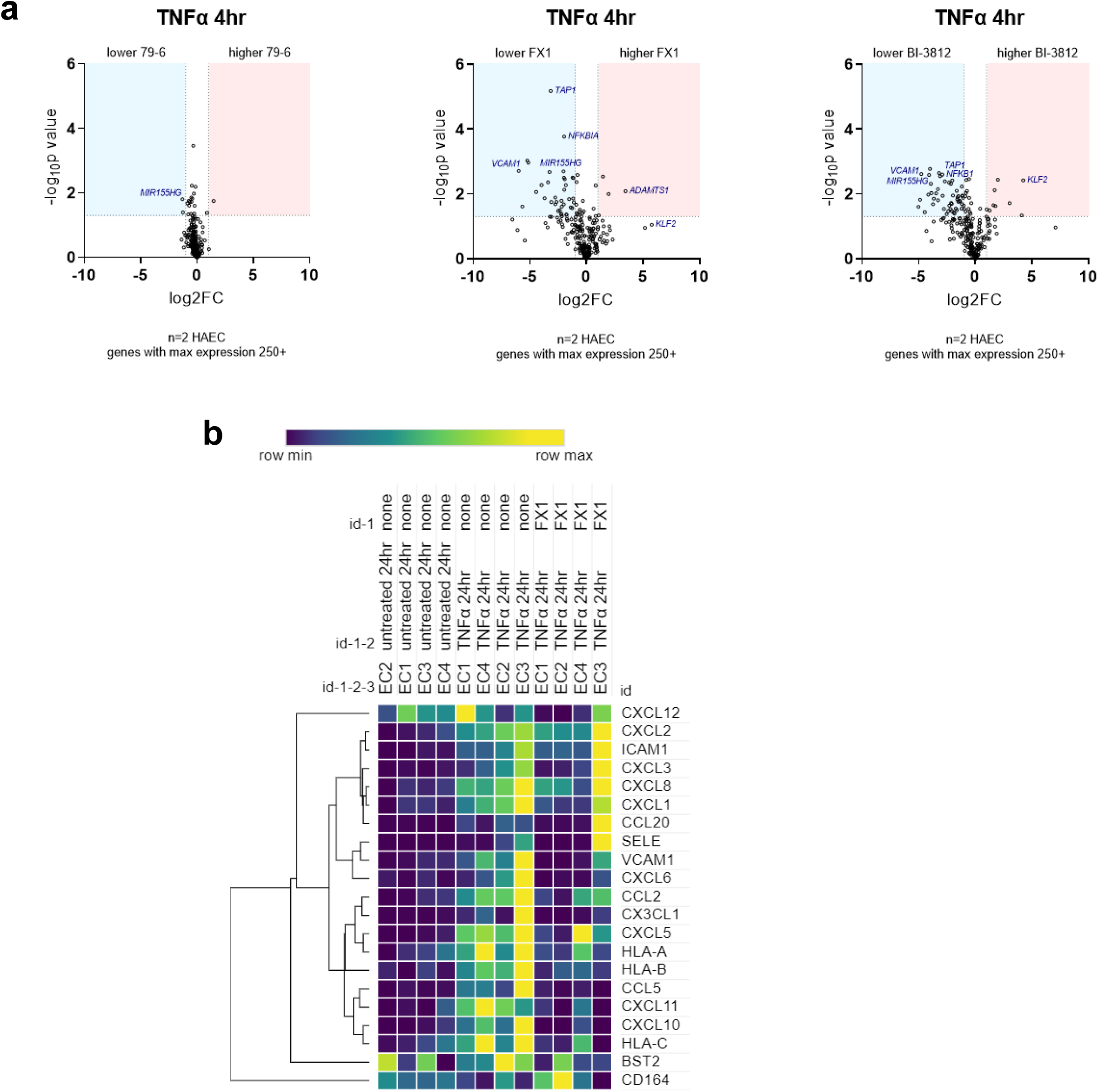
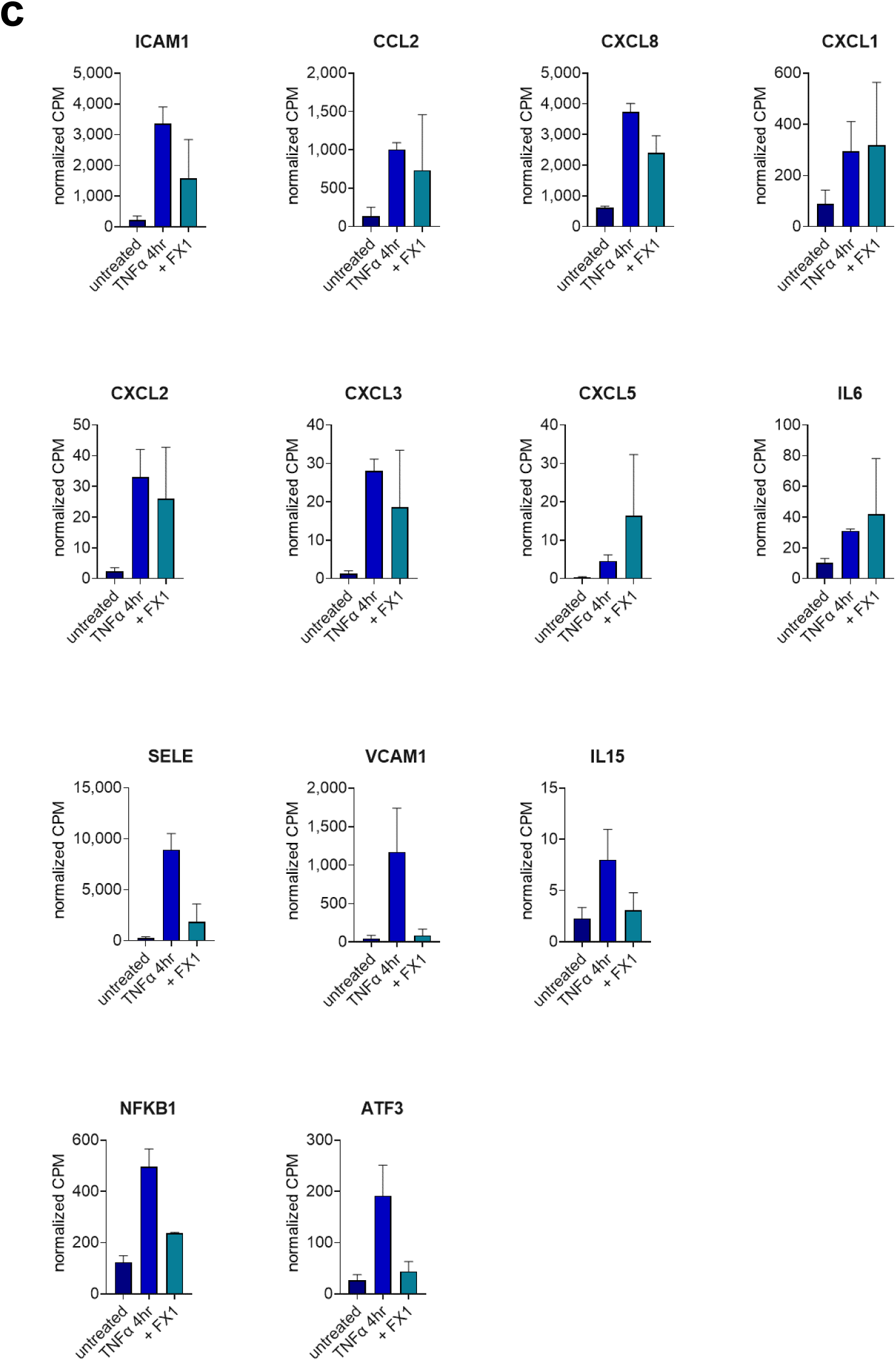
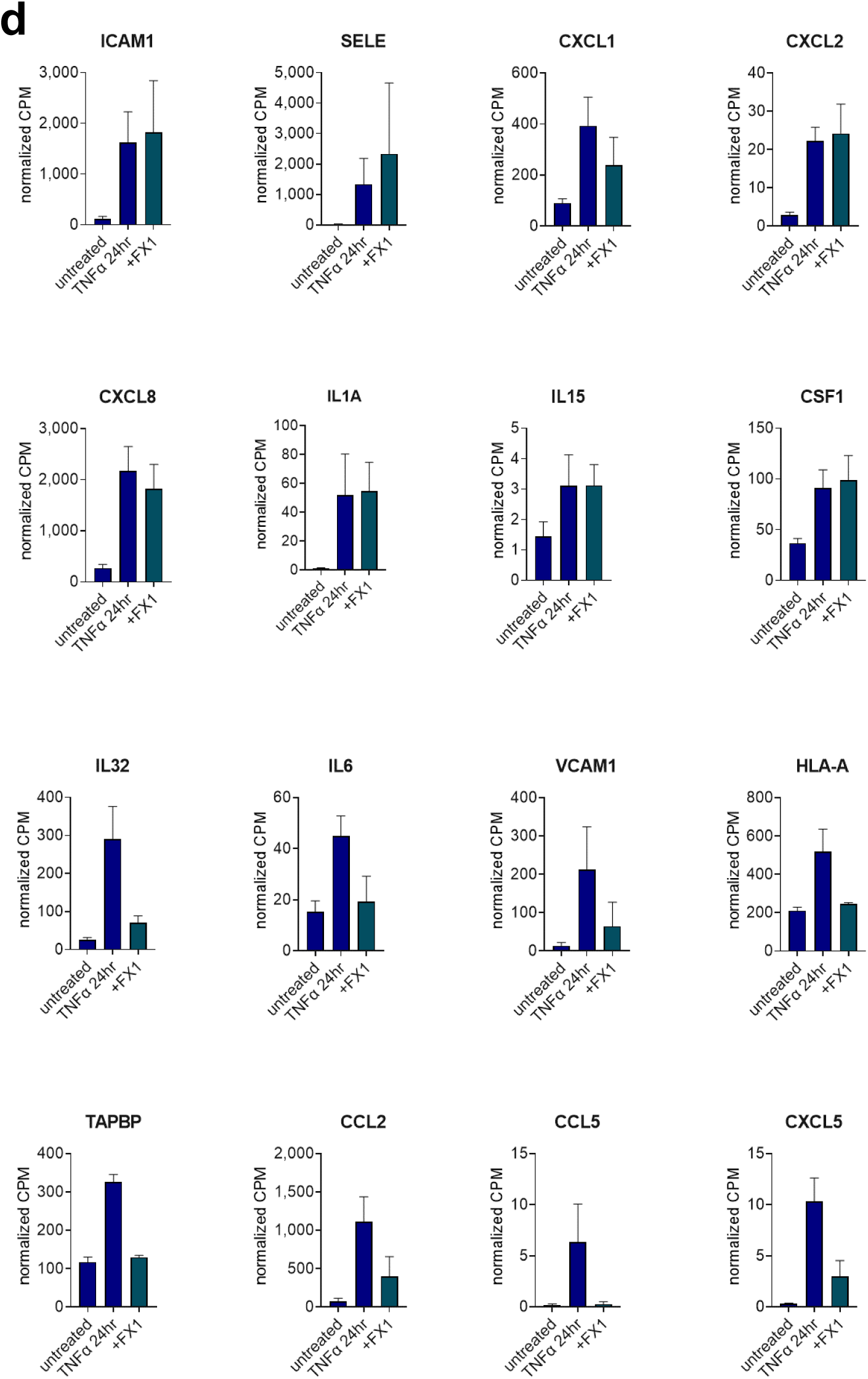
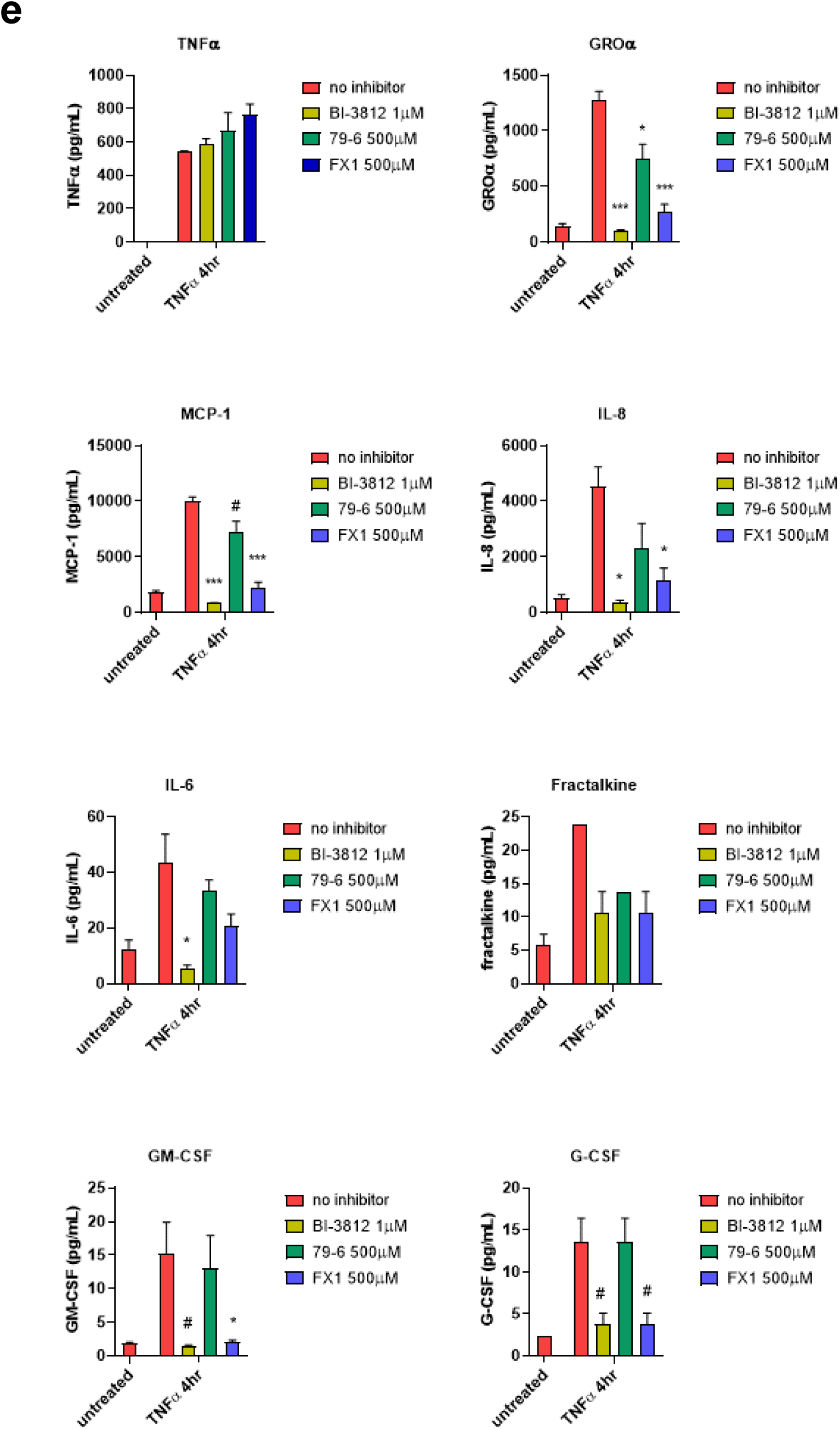
(a) Volcano plots from Nanostring comparing gene expression in TNFα-treated HAEC in the absence or presence of 79-6, FX1 or BI-3812. (b) Heat map from RNA-Seq results of primary HAEC treated with TNFα for 24hr, in the absence or presence of FX1 (n=4 biological replicates). BTB domain inhibition affects endothelial pro-adhesive gene and protein expression. (c) Graphs show average normalized CPM counts for selected inflammatory genes. BTB domain inhibition affects endothelial pro-adhesive gene and protein expression. (d) Graphs show average normalized CPM counts for selected inflammatory genes.

**Supplemental Figure 8.**
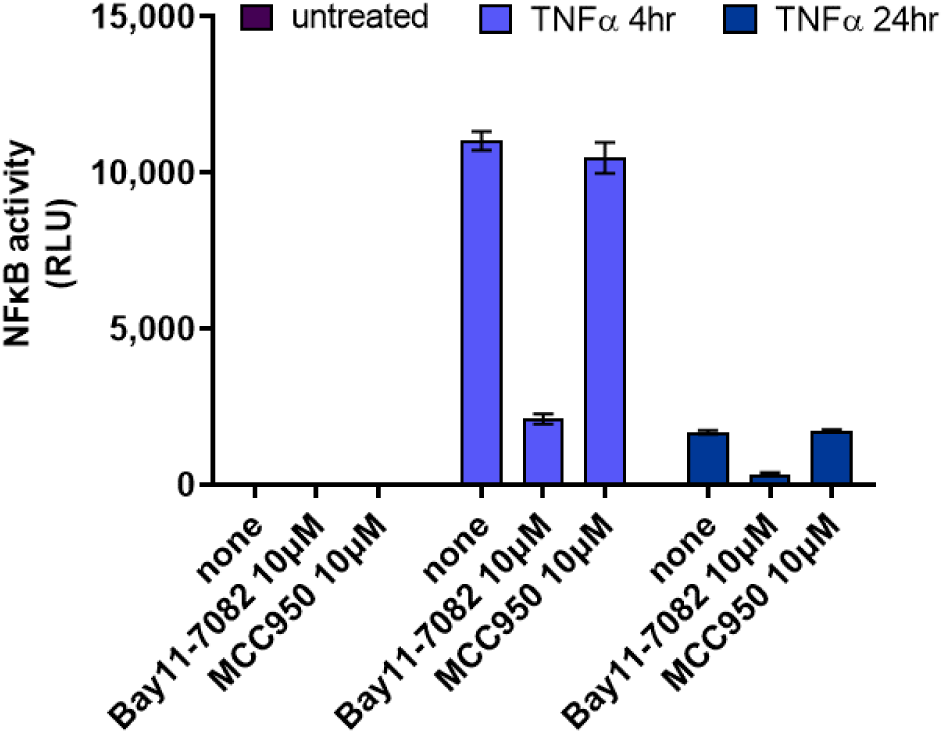
NFKB-TIME endothelial cells were stimulated with TNFα for 4hr or 24hr, in the absence or presence of the NFKB/NLRP3 inhibitor Bay11-7082, or the NLRP3 inhibitor MCC950 as a control for specificity. NFKB transcriptional activity was measured by luciferase assay.

**Supplemental Figure 9.**
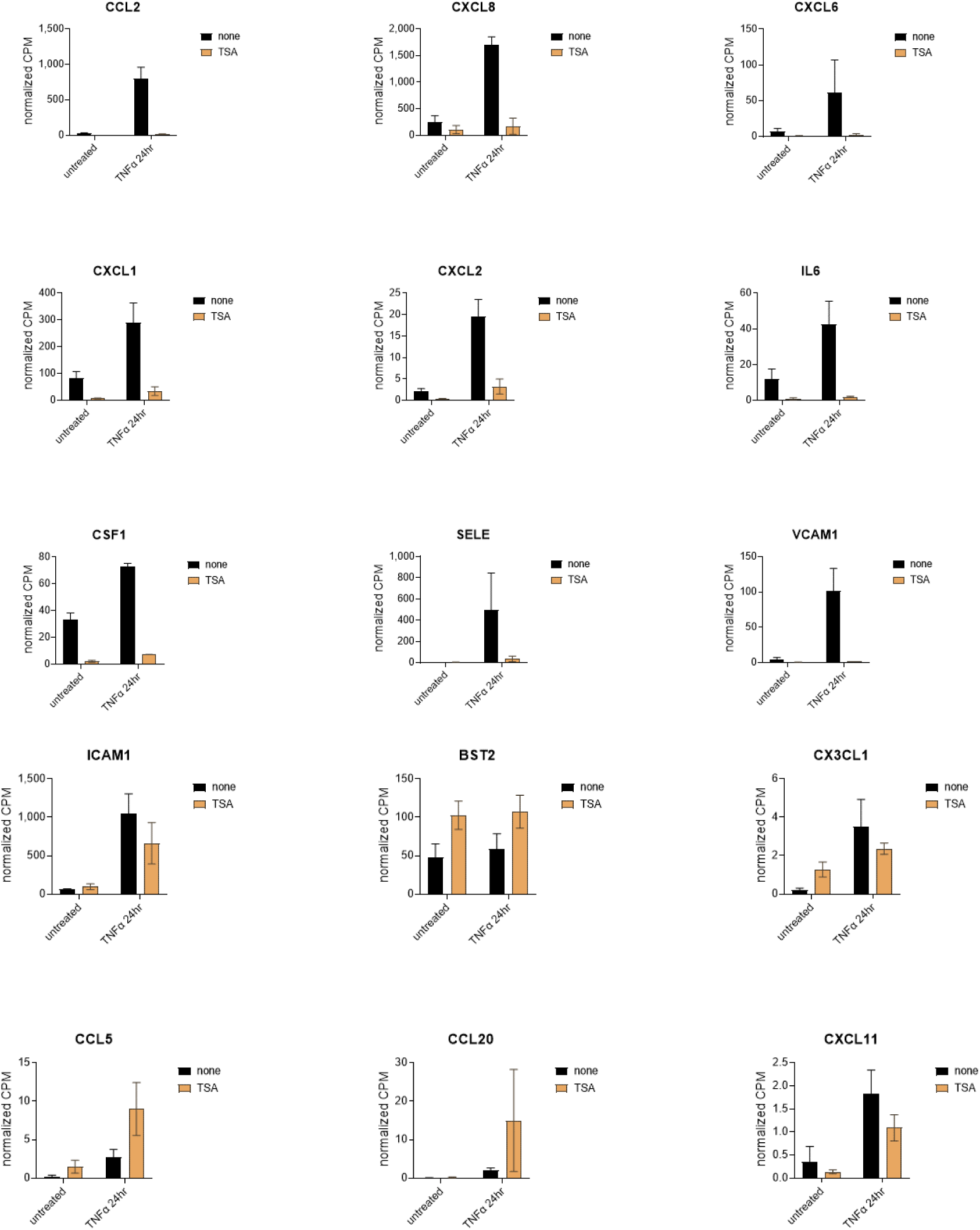
HDACs and endothelial pro-adhesive activation. Primary HAEC (n=2 biological replicates) were treated with the HDAC inhibitor TSA and stimulated with TNFα. Gene expression was measured by RNA-Seq. Graphs show expression of TNFα-induced genes.

## Notes

### Competing Interest Statement

The authors have declared no competing interest.

